# FlyTomo: A streamlined software for on-the-fly cryo-ET data processing and diagnosis

**DOI:** 10.1101/2025.07.24.666517

**Authors:** Zheyuan Zhang, Cheng Peng, Weiping Zhang, Jiaming Liang, Kexin Liu, Yong Chen, Junxia Zhang, Rui Liang, Yutong Song, Sai Li

**Author notes:** These authors contributed equally (Zheyuan Zhang, Cheng Peng, Weiping Zhang).

## Abstract

Cryo-electron tomography (Cryo-ET) combined with subtomogram averaging (STA) enables the structural elucidation of macromolecular assemblies in their native environments. However, the widespread adoption of cryo-ET has been limited by its labor-intensive, expertise-dependent data processing workflow. Here we present FlyTomo, a software that streamlines data processing from frame alignment to STA with high-throughput for authentic cryo-ET scenarios. During data acquisition, FlyTomo performs real-time diagnosis, enabling prompt feedback on sample quality, microscope performance and structural features. After acquisition, it aggregates diagnostic metrics into an interactive overview, guiding users through data review and refinement. FlyTomo also curates raw and processed data into structured directories to simplify data management and archiving. We validated FlyTomo across a diverse set of authentic cryo-ET samples, including purified enveloped viruses and cryo-lamellae, on multiple microscopes and cameras, achieving structures at resolutions ranging from 3.4 to 5.7 Å. Collectively, by integrating accuracy, scalability and usability, FlyTomo reduces the technical barrier for in situ structural biology using cryo-ET.

## Introduction

Recent advancements in cryo-electron tomography (cryo-ET) and subtomogram averaging (STA) have led to a surge of biological discoveries resolved from native contexts^1,2^. However, despite these breakthroughs, cryo-ET remains a specialized technique with limited adoption in structural biology. According to the electron microscopy data bank (EMDB) depositions over the past decade, cryo-ET, including STA and tomogram datasets, contributed only approximately 7-12% of all cryo-EM submissions in the last five years, down from 15-24% in the preceding five years. Compared to cryo-EM single-particle analysis (SPA), the dominant cryo-EM approach, broad applicability of cryo-ET is hampered by its low-throughput, low-resolution and long learning curve.

Cryo-ET poses exceptional challenges compared to SPA. The need to collect tilt-series (TS) data under low-dose conditions, while continuously tilting the specimen stage, places stringent requirements on microscope alignment and stability. Monitoring of tilt-axis offset^3,4^ and coma^3,5,6^ is required throughout the acquisition sessions to maintain alignment fidelity. Furthermore, cryo-ET targets, including isolated complexes^7^, enveloped viruses^8,9^ and cells^10^, are often structurally heterogeneous, scarce in abundance, and embedded in crowded biological contexts. These complexities often necessitate volumetric inspection of areas of interest (AOIs) to assess structural features and distribution of the proteins of interest (POIs), which cannot be reliably inferred from single micrographs alone^11^. Ideally, diagnostic evaluation of microscope performance, data quality, and sample integrity should occur on-the-fly during TS acquisition, enabling corrective measures during a live session. However, in conventional workflows, the significant latency between data collection and analysis precludes timely feedback.

Post-acquisition, terabytes of raw TS data must undergo an extensive multistage processing workflow before high-resolution structures can be resolved^1,12–14^. This briefly includes five main steps categorized into a pre-processing stage: (1) frame alignment to correct beam-induced motion, and contrast transfer function (CTF) estimation and correction, (2) TS assembly and alignment, (3) tomogram reconstruction and denoising for identification of POIs; followed by a structural determination stage: (4) POIs identification, STA and classification; and eventually a post-processing stage: (5) local refinements of CTF and TS alignment, map sharpening, and model building. Each step involves benchmarking various tools and parameter sets, along with inter-software format conversion. Additionally, the increasing volume of cryo-ET datasets^15^ imposes significant burdens on data transfer, storage, and computational resources. Consequently, workflows vary widely across laboratories depending on the biological system and user expertise. The lack of standardization, coupled with a steep learning curve, presents a major obstacle for non-specialist users and impedes broader adoption of cryo-ET.

To address these challenges, we introduce FlyTomo, a robust, user-centric, and integrated software platform designed to streamline and democratize cryo-ET data processing. FlyTomo contains three key innovations: (1) A cryo-ET workflow covering frame alignment through to STA with empirically optimized parameters. Its flexible architecture supports diverse microscope systems and data acquisition schemes to accommodate heterogeneous experimental requirements; (2) An on-the-fly feedback system, providing rapid tomogram reconstruction and diagnostic metrics within minutes post-acquisition to facilitate rapid assessment of sample quality, instrument stability, and data integrity; and (3) An offline intervention-reprocessing loop that provides a convenient intervention function for refinement of individual steps based on its diagnosis on the data processing workflow. These innovations were based on its efficient computational strategies, including multi-CPU/GPU parallelization, data compression, and in-memory processing. FlyTomo has been extensively tested on real-world cryo-ET scenarios, achieving resolutions in the range of 3.4 to 5.7 Å. Overall, FlyTomo significantly reduces technical barriers, enabling researchers with varying levels of expertise to achieve high-resolution in situ structures with greater throughput and confidence.

## Results

### Overview of FlyTomo

FlyTomo streamlines cryo-ET data processing with high throughput and resolution. It’s compatible with both conventional TS collection methods, which record AOIs individually using continuous, bi-directional or dose-symmetric^3^ schemes, and beam-image shift (BIS) TS collection method which records multiple AOIs simultaneously. Once initiated by a click, FlyTomo continuously monitors the data output and automatically processes the raw movie data.

The standard preprocessing includes motion correction by MotionCor2^16^, CTF estimation by gctf^17^, TS curation (dark image deletion, dose weighting and assembly of TS), TS alignment by IMOD^18^ or AreTomo^19^, tomogram reconstruction, deconvolution and snapshotting. For a TS, these procedures are completed within minutes after all tilts have been recorded, generating a denoised tomogram for rapid volumetric inspection of the sample. Next, five optional functions, including 3D-CTF correction by NovaCTF^20^, denoising by IsoNet^21^, membrane segmentation by MemBrain-Seg^22^, air-water interface (AWI) detection using our algorithms, are available for tomogram processing. The 3D-CTF corrected tomograms facilitate STA or denoising, while denoised tomograms enhance POI identification. Membrane segmentation supports identification and initial orientation assignment for membrane-bound POIs, whereas AWI detection module is optimized for cellular tomography applications.

Throughout the processing pipeline, intermediate outputs indicative of microscope performance, sample integrity, and structural quality are promptly analyzed, diagnosed, visualized, and communicated to users for immediate assessment. Following coarse TS alignment, FlyTomo provides a GUI that enables manual inspection and refinement of images flagged by the diagnostic module as potentially problematic. Users may iteratively reconstruct tomograms following removal or realignment of suboptimal data until satisfactory quality is achieved, thereby converting the traditionally labor-intensive TS alignment into an efficient, diagnosis-driven workflow. These procedures are followed by a structural determination module, consisting of POI identification and STA. FlyTomo executes particle picking using pytom-match-pick^23^, subsequently cropping subtomograms and preparing the dataset for automated STA in Dynamo^24^, executed via a structured parameter input file without manual GUI activation (Fig. 1a). FlyTomo features a user-friendly GUI optimized for effective interaction and structured graphical feedback (Fig. 1b).

**Fig. 1.**
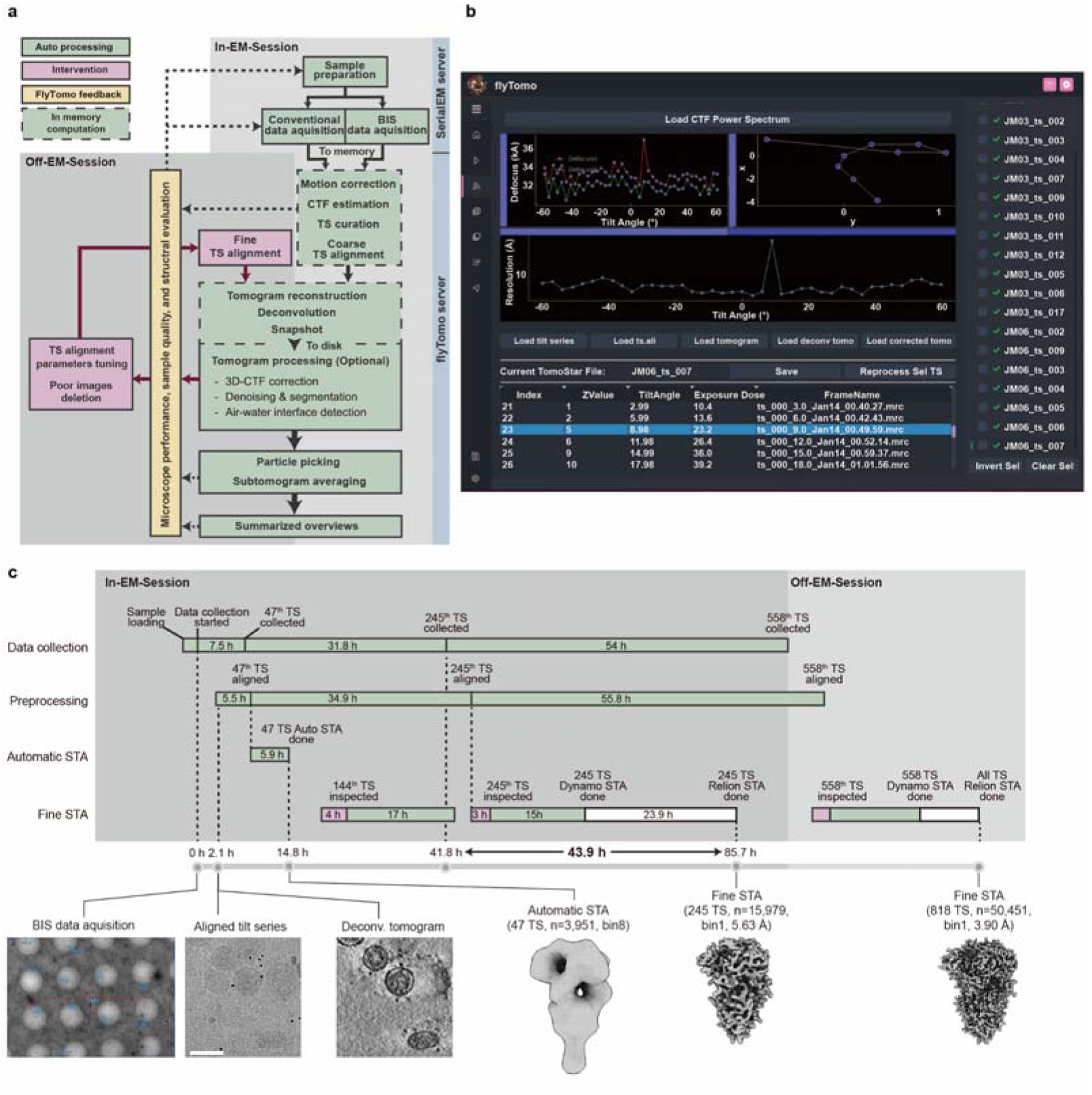
Introduction to the workflow, interface and execution of FlyTomo. **a**, Flowchart illustrating FlyTomo streamline. The green boxes are automatic steps with some can optionally computed in memory for acceleration (outlined with dotted line), the magenta boxes involve steps that need human intervention, and the yellow boxes are the visualized and statistical feedback on the sample, data, and microscope. **b**, The GUI of a typical page in FlyTomo. Each set of TS data is listed in the right panel. When selecting a certain set of data, information of all tilt images is showcased in the lower panel with preprocessing results in the upper panel. **c**, A timeline of the on-the-fly processing on HCoV-229E using FlyTomo. Scale bar: 100 nm. Color coding follows **a**, except white for external software.

### On-the-fly data analysis and in-memory computation

FlyTomo’s on-the-fly data analysis benefited from its automated data processing, in-memory computation and data compression functions. To facilitate automation, FlyTomo implements two empirical protocols optimized for specific cryo-ET scenarios. The molecular tomography protocol integrates empirically selected algorithms and parameters suitable for cryo-ET data comprising isolated macromolecular complexes, typically characterized by fiducial markers, carbon films, relatively thin ice, and clean backgrounds. In contrast, the cellular tomography protocol is specifically optimized for cryo-focused ion beam (FIB)-milled lamella data, which generally lack fiducials and feature thicker ice, crowded backgrounds, and pretilt angles. By supplementing a pre-defined Dynamo STA parameter form and a template for POI matching, FlyTomo is capable of automatically processing raw movies to structures.

Conventional cryo-ET workflows involve repetitive read-write operations on storage disks between processing stages, generating substantial I/O demand. To mitigate intensive storage and network demands, FlyTomo contains an in-memory computation function, converting the computational environment for data processing steps from hard drive to memory. Using this approach, FlyTomo directly reads raw movie data from microscope servers into memory on processing servers, subsequently executing in-memory from motion correction to taking snapshots. Upon completion, intermediate results are either discarded from memory or transferred to designated storage disks. This function reduces server I/O intensity by approximately 60% for processing MRC-format movie files (Supplementary Table 1). Additionally, FlyTomo efficiently manages storage requirements for intermediate results and raw data, typically extensive in number and size, by retaining only essential files for tomogram reconstruction and STA within systematically categorized directories (Supplementary Fig. 1). Non-reproducible raw data, such as movies of each tilt, undergo parallelized lossless compression, achieving over 90% reduction in storage usage for MRC-format raw data.

FlyTomo also supports BIS TS acquisition schemes and incorporates automated parallel processing leveraging multiple CPUs and GPUs (Supplementary Fig. 2). Tasks are dynamically allocated to available hardware resources based on user-defined configurations, enabling concurrent execution and optimal hardware utilization. Collectively, these enhancements substantially improve throughput, simplify data sharing, storage, and management, and enable researchers to concentrate on analysis rather than protracted computational processes, particularly beneficial for high-resolution, large-scale cryo-ET projects (Supplementary Table 1).

### In-EM-Session and Off-EM-Session workflows

To further improve throughput, we next sought to improve data acquisition efficiency while ensuring optimized data quality. This is achieved by designing an In-EM-Session workflow, when users are in an active cryo-ET data acquisition session; and an Off-EM-Session workflow, when the microscope session has terminated. During the In-EM-Session, the workflow prioritizes automation, throughput, and immediate diagnostic feedback due to intensive microscope operation procedures, including microscope setup, grid and AOI screening, and TS acquisition software configuration. This approach allows users to benefit from real-time diagnostics related to microscope performance, data quality at various processing stages, specimen ice profile, tomographic sample overview, and POI structural assessment. Users can promptly address any deficiencies identified by FlyTomo or user-defined criteria by adjusting microscope settings, altering imaging areas, or switching grids during the session.

The Off-EM-Session workflow emphasizes refining specific processing steps that require manual intervention and large-scale computational resources to resolve high-resolution structures. Revealed by the diagnostic feedback, FlyTomo proactively alerts users to potentially problematic datasets and provides an interactive interface for detailed data review and refinement. Users can iteratively reprocess updated datasets to STA using a continuing function within FlyTomo. For low-binning, full-dataset processing requiring extensive computation, FlyTomo offers a one-click functionality that reformats results for compatibility with RELION4/5^25,26^, facilitating resolution refinement.

To quantitatively evaluate throughput and validate FlyTomo’s practicality in an authentic cryo-ET scenario, we applied it to a molecular tomography project conducted over a four-day microscope session. The project aims at determining a near-atomic resolution structure of the spike (S) protein from isolated, active human coronavirus 229E (HCoV-229E). Previous experiments indicated the structural integrity and abundance of HCoV-229E S are sensitive to propagation, purification, and storage conditions. Rapid structural evaluation of S, otherwise unclear from deconvoluted tomograms, was therefore essential. By integrating FlyTomo’s In-EM-Session workflow with PACEtomo^4^, we collected 47 TS within 7.5 hours of initiating data acquisition. Within 1.4 hours post-acquisition, these TS underwent alignment, reconstruction, and deconvolution to produce denoised tomograms. Utilizing a pre-trained model for S protein identification in ilastik^27^ and predefined STA parameters derived from earlier tests, FlyTomo automatically resolved an initial S protein structure within 14.8 hours. Although derived from 8 × binned tomograms, the map sufficiently demonstrated that the majority of S proteins existed in prefusion conformation and remained structurally intact on HCoV-229E (Fig. 1c).

With confidence in the sample quality established, we proceeded with data collection and utilized the Off-EM-Session workflow for the inspection and correction of suboptimal data when the microscope was collecting TS. At 85.7 hours, a 5.63 Å resolution structure was refined from 245 TS. Eventually, combining the 558 TS collected during this session with an additional 260 collected previously, a 3.90 Å resolution structure was resolved (Fig. 1c, Supplementary Table 2). Overall, this evaluation demonstrated that FlyTomo significantly enhances cryo-ET data processing efficiency, optimizes data quality during active microscope sessions, and effectively identifies and refines suboptimal data to facilitate high-resolution structure determination.

### On-the-fly diagnosis

FlyTomo incorporates an on-the-fly diagnostic function that transforms essential intermediate results indicative of microscope status, data quality, and sample integrity into actionable visualizations and summaries. These diagnostics are provided through various interactive GUI panels (Fig. 1b) and promptly generated comprehensive summaries.

FlyTomo’s diagnostic module comprises four key functions. First, it addresses abnormal motion caused by factors such as electron beam instability, stage movements, or sample defects near the AOI, which can lead to unsuccessful motion correction and blurred averaged tilt images. These images often abrogate a TS if concatenated into it. FlyTomo facilitates the identification and removal of these problematic images by visualizing defocus values, CTF-fitting resolution, and power spectra of each tilt image post-CTF estimation. Outlier measurements can be rapidly distinguished and excluded (Fig. 2a). Second, the TS alignment diagnostic assesses microscope stability. FlyTomo plots fiducial or patch-tracking trajectories alongside residual mean errors and standard deviations, helping to identify misplaced fiducials, mismatched patches, intrinsic motion in TS, and improper beam tilt calibrations (Fig. 2b, c, Supplementary Fig. 3). These identified TS can be promptly corrected or discarded through IMOD within FlyTomo’s GUI. Third, FlyTomo monitors and visualizes shifts of AOIs during TS collection, particularly at higher tilt angles. Overlapping fields of view are displayed, highlighting substantial shifts for user evaluation (Fig. 2d). Additionally, FlyTomo continuously summarizes beam tilt information from fiducial trajectories to monitor the microscope performance^28^ and provides tomogram snapshots for comprehensive sample inspection. Lastly, the assembly, distribution and structural integrity of POIs are often only assessable in tomograms or even preliminary structures for actual cryo-ET projects. To allow prompt volumetric evaluation of the sample, FlyTomo prepares aligned TS and reconstructs deconvoluted tomograms within minutes, which are accessible from the GUI.

**Fig. 2.**
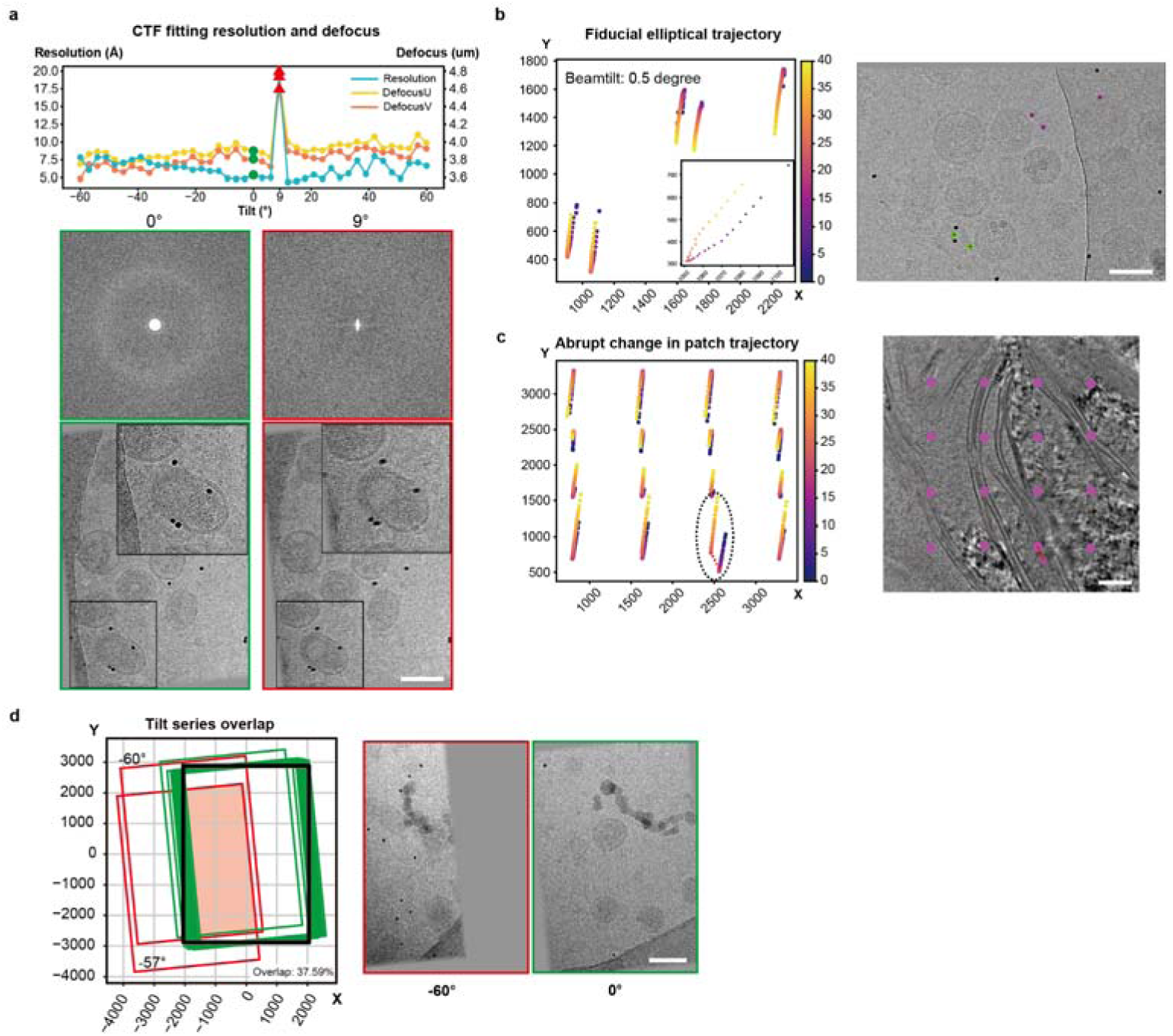
FlyTomo feedback for data quality and processing performance assessment. **a.** The defocus values (DefocusU and DefocusV) and fitted resolutions across a TS (top) with images and corresponding power spectra at 0° and 9°, respectively. **b.** Fiducial track analysis. The fiducial track with high beam tilt. **c.** Patch track analysis. The abrupt change in patch trajectory. **d.** Overlapping regions of all images in the TS. The orange area represents the common regions shared across different tilt angles; the rectangles indicate the spatial distribution of individual images at each tilt angle. Red, large image shifts; Green, subtle image shifts; Black, reference view area. Scale bar: 100 nm.

### Cryo-lamellae air-water interface detection

Accurate measurement of ice geometry is a necessary step in cryo-ET. First, tomogram reconstruction requires ice thickness and center estimations to define boundaries between signal-containing and void volumes. Second, certain TS alignment algorithms, such as AreTomo, relies on accurate ice thickness measurements. For instance, mistakenly estimated ice thickness in AreTomo will deteriorate its TS alignment, as assessed from both the ribosome signals in the x-y plane and platinum deposition in the x-z plane (Fig. 3a). Third, reliable ice geometry measurements enable accurate determination of lamella pretilt angles. However, measuring ice geometry in cryo-lamellae from vitrified cells remains challenging due to intrinsic pretilt angles and uneven ice thickness, often necessitating labor-intensive manual measurements.

**Fig. 3.**
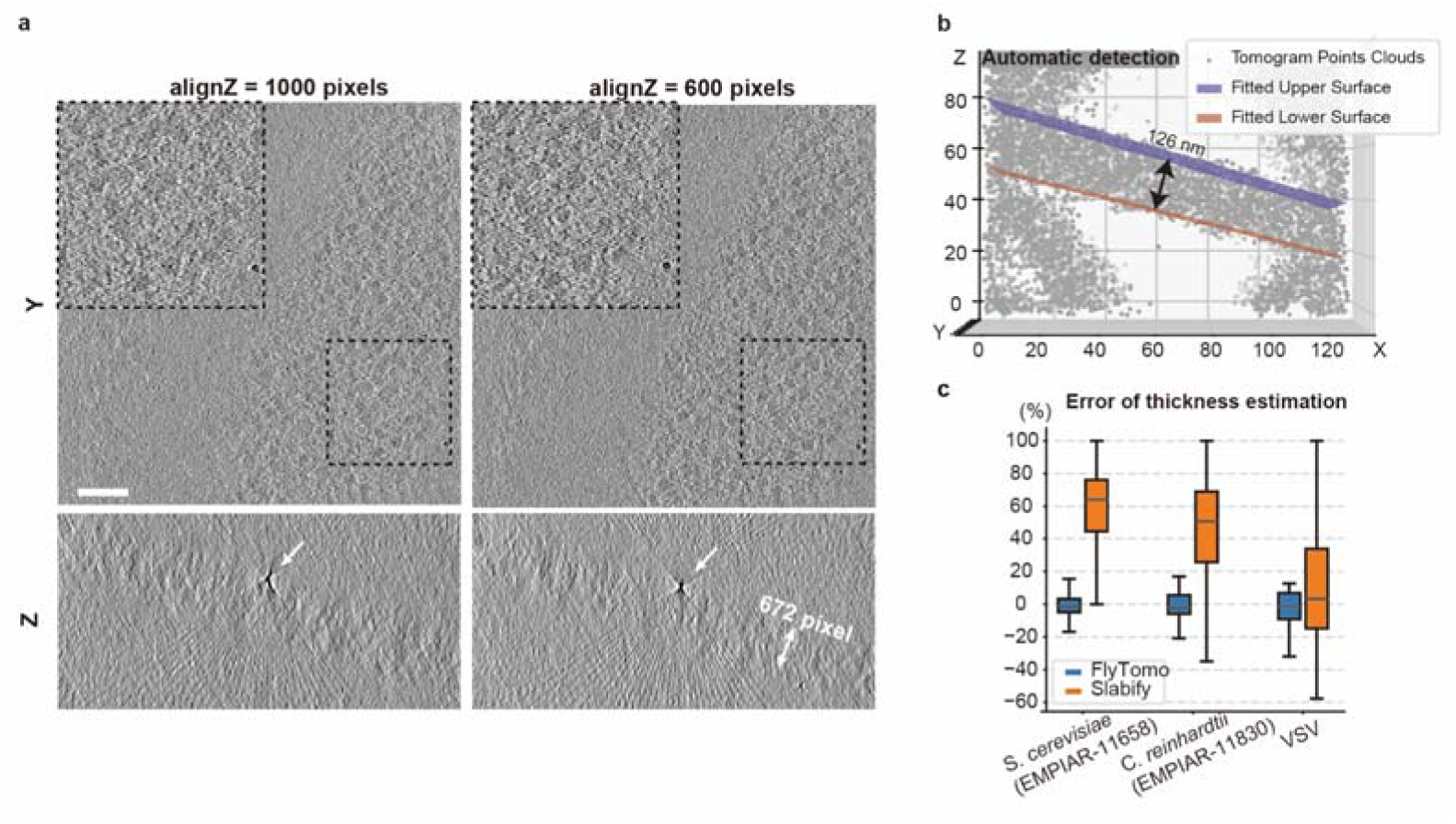
Detection of AWI in FlyTomo. **a**, Effect of correct alignZ parameter in AreTomo on alignment quality. Same TS aligned with -alignZ = 1000 (left) vs. -alignZ = 600 (right); true thickness: 672 pixels. **b**, The AWI detection result of an example tomogram. **c**, Comparison of percentage error of lamellae thickness estimation between FlyTomo (blue) and Slabify (orange), outliers excluded.

To automate this process, we introduced a Shannon entropy-based algorithm for automatic detection of AWIs within tomograms, simplifying the geometry model by assuming planar interfaces. The algorithm involves filtering the tomogram with an optional 3D SIRT-like filter to enhance contrast, calculating Shannon entropy to evaluate local randomness, segmenting the ice layer via Otsu’s thresholding and morphological closing, and fitting planes to the interfaces using a RANSAC algorithm (Fig. 3b).

FlyTomo uses these detected interfaces to estimate lamella thickness, subsequently rendering thickness profiles for selected datasets (Fig. 4f). To validate our automated approach, we tested it on three distinct datasets containing lamellae from *S. cerevisiae* (EMPIAR-11658, n = 270), *C. reinhardii* (EMPIAR-11830, n = 59), and cells infected with vesicular stomatitis virus (VSV, this study, n = 23). For benchmarking, these data were also manually measured as ground truth, alongside automated measurements by Slabify^29^. Our method demonstrated greater accuracy with narrower interquartile ranges and smaller median errors across all datasets (Fig. 3c). Additionally, the identified AWIs facilitated automatic calculation of lamella pretilt angles (Fig. 4e). Algorithm parameters are adjustable for specific tomograms, yet our method generally remains robust across entire datasets (Supplementary Fig. 4). Results, including input parameters, calculated lamella thickness, pretilt angles, and fitted plane equation parameters, are exported as spreadsheets. The exported parameters enable users to generate accurate masks for downstream analyses.

**Fig. 4.**
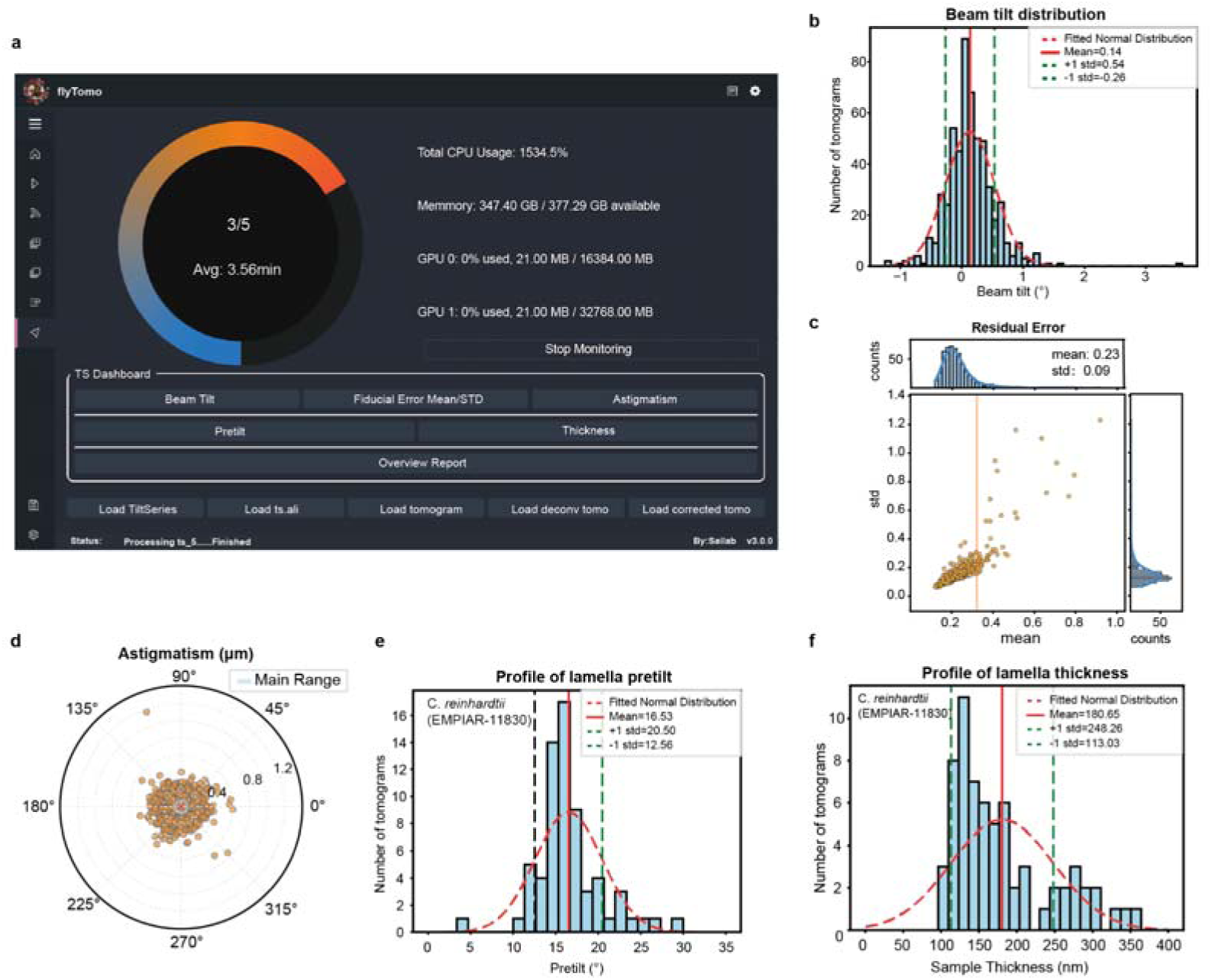
Summarized overview of the processed dataset in FlyTomo. **a**, The FlyTomo GUI provides a real-time overview of processing status and data quality metrics. **b**, Histogram of beam tilt across collected tomograms. **c**, The mean and STD of residual errors across the TS with fiducial alignment. The orange line represents the mean residual error of 0.35. **d**, Polar plot of astigmatism magnitude and orientation. **e,** Histogram of lamella pretilt angles. **f**, Histogram of lamella thickness values for a cryo-FIB dataset (EMPIAR-11830).

### Summarized overview

Efficient curation and quality control for the streamline and the subsequent large datasets is essential for high resolution and throughput cryo-ET. To address this requirement, FlyTomo includes a comprehensive summarized overview function, designed to facilitate rapid data evaluation and quality assessment at a glance (Fig. 4a). This function systematically aggregates diagnostic metrics for each TS into a customizable summary report, enabling quantitative evaluation of acquisition quality. Specifically, it computes and collates statistics for essential parameters such as beam tilt (Fig. 4b), residual mean error resulting from TS alignment (Fig. 4c), astigmatism (Fig. 4d), lamella pretilt (Fig. 4e), and estimated lamella thickness (Fig. 4f).

Additionally, FlyTomo generates representative snapshots from the central slices of corresponding deconvolved tomograms, offering immediate visual impression on reconstructed volume quality. These snapshots quickly inform users about the specimen integrity, ice contamination levels, and spatial distribution of POIs. All statistical data and visual snapshots are combined into an editable document, creating a succinct and navigable report.

Together, the summarized overview significantly enhances the on-the-fly diagnostic function, streamlines data curation, and allows researchers to efficiently monitor the quantity and quality of TS acquired during the In-EM-Session. It also helps users to identify and exclude problematic data and prioritize high-quality TS for subsequent processing during the Off-EM-Session, thus streamlining the post-acquisition workflows.

### Adaptivity to user preferences and microscope systems

Given the diverse nature of cryo-ET samples, FlyTomo is designed to be highly adaptable, allowing integration into a wide range of data processing workflows, customization according to user preferences, and compatibility across multiple microscope and camera systems. FlyTomo can support real-time cryo-ET sessions on-site and accommodate standalone import of raw movies in various formats, including MRC, TIFF, EER, and BZ2. Processed images are exported in MRC format, while all associated metadata, such as tilt angles and electron dose, are systematically saved in STAR files, a format recognized by the EM community for its scalability and machine readability.

FlyTomo allows flexibility at each stage of the processing pipeline, enabling users to skip specific steps as necessary. We also made FlyTomo adaptable to various microscope and camera configurations, including a Titan Krios equipped with a Gatan K3 camera, and a Titan Krios G4 with a Falcon 4i camera, yielding resolutions from 3.4 to 5.7 Å. Comprehensive testing across these authentic cryo-ET scenarios confirms FlyTomo’s adaptability, performance, throughput efficiency, and achievable resolution.

FlyTomo has been primarily tested on authentic cryo-ET scenarios across multiple microscope systems. We categorized the scenarios into molecular and cellular tomography, selecting seven representative samples to showcase the software’s applications. Most of these samples were prepared in-house, and extensive datasets were acquired on-site to rigorously test FlyTomo’s In-EM-Session diagnostic capabilities. The test samples included purified 20S proteasome as a standard control, isolated viruses such as HCoV-229E, Influenza A Virus (IAV), VSV for molecular tomography, and ribosomes from cryo-lamellae of *S*. *cerevisiae* (EMPIAR-11658) for cellular tomography. An overview and detailed characteristics of these test samples are summarized (Table 1), with additional specifics on data collection and processing methods outlined (Supplementary Table 2). For the majority of these samples, FlyTomo was applied during the In-EM-Session, processing data simultaneously with the data acquisition to test its on-the-fly diagnosis.

**Table 1.** Samples tested by FlyTomo.

| Type | Species/Strain | Molecule | Mass (kDa) | Fiducials | Microscope | Particle feature | Motivation |
| --- | --- | --- | --- | --- | --- | --- | --- |
| Purified protein | *T. acidophilum* | Proteasome | 720 | No | Titan Krios, Gatan K3, stable | D7, homogeneous | Proof-of-principle |
| Flexibly assembled macromolecules | HCoV-229E | S | 600 | Yes | Titan Krios, Gatan K3, stable | C3, spread on membrane | High resolution |
|  | IAV | HA | 190 | Yes | Titan Krios G4, Falcon 4i, stable | C3, spread on membrane | High resolution |
|  | VSV | RNP | 100 (subunit) | Yes | Titan Krios, Gatan K3, stable | Helical lattice within membrane | High resolution |
| Cell lamellae | *S. cerevisiae* | 80S Ribosome | 3100 | No | Titan Krios G4, Falcon 4i, stable | C1, regional | Dealing with crowded cell environment |

### Testing FlyTomo on standard and molecular tomography samples

To assess the performance and general applicability, we tested FlyTomo on diverse samples including purified complexes and intact enveloped viruses. These evaluations focused on three metrics: diagnostic effectiveness, processing throughput, and achievable resolution. We first validated FlyTomo’s performance on purified proteasome to establish proof of principle. FlyTomo processed 40 fiducial-less TS and identified 34,380 particles, achieving a final resolution of 3.4 Å (Fig. 5a). The resolution was evidenced by side-chain densities. We confirm that for benchmark samples, even in absence of fiducials, the FlyTomo-processed data contain signals for near-atomic resolution structure determination.

**Fig. 5.**
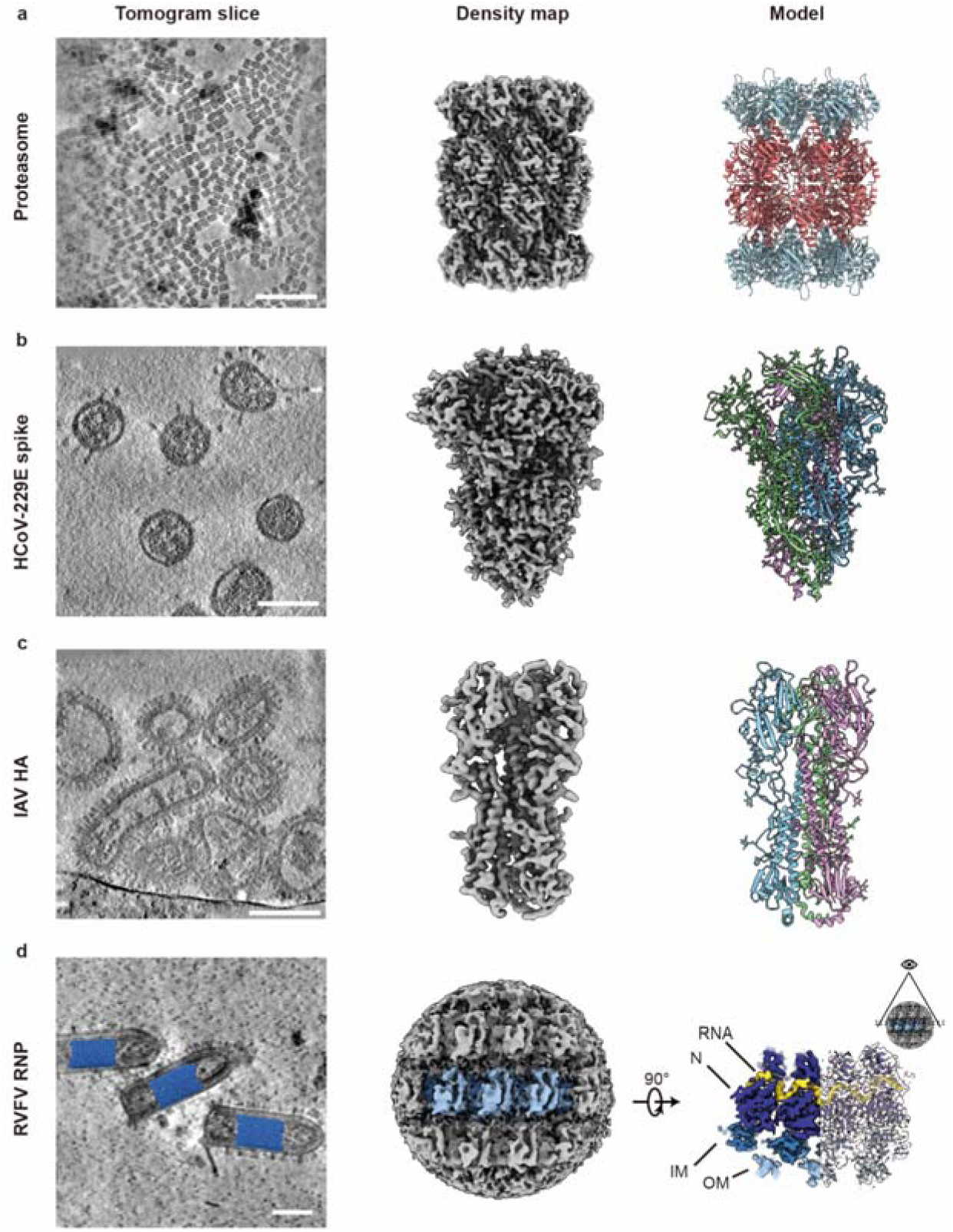
FlyTomo processing on purified macromolecule samples. **a.** Proteosome tomogram slice (left), STA density map (center), and corresponding atomic model (right). **b.** HCoV-229E tomogram slice (left), STA density map of S-trimer (center) and corresponding atomic model (right). **c.** IAV tomogram slice (left), STA density map of HA (center) and corresponding atomic model (right). **d.** VSV trunk reconstructions shown in tomogram slice (left). The STA density map (right) fitted atomic model (PDB: 7UWS). Scale bars, 100 nm.

We next evaluated FlyTomo on representative molecular-tomography specimens. Cryo-ET is frequently used to study assembly and membrane-fusion mechanisms of heterogeneous enveloped viruses. Here, our goal was to obtain high-resolution structures of the HCoV-229E spike trimer, IAV hemagglutinin (HA) trimer, and the VSV ribonucleoprotein (RNP).

On HCoV-229E, spike trimers are sparse and structurally heterogeneous (Fig. 5b). Preliminary cryo-EM screening indicated that each virion harbors approximately 9 conformationally stable spikes on average, complicating high-resolution analysis. From 818 tomograms (5,621 virions), a 3.90 Å spike map was determined, with local resolution extending to 3.75 Å (Fig. 5b; Supplementary Fig. 5). The resulting atomic model aligns with the recombinant “tight” conformation (PDB:7CYC)^30^. This example demonstrates that low-abundance, heterogeneous targets benefit from FlyTomo’s streamlined, high-throughput pipeline.

HA is a 200 kDa trimer residing on the IAV envelope. As a recombinant protein, HA adopts severe preferred orientations in vitreous ice and imposes challenges for SPA^31^. Its abundant presence on viral surface (Fig. 5c) makes structural determination by SPA from the perimeter of the viral envelope challenging^32^. Previous cryo-ET studies achieved only 8.9 Å resolution for on-virion HA^33^, likely limited by its modest molecular weight. Using 65 tomograms (363 virions), FlyTomo resolved a 3.6 Å HA trimer map with local resolution up to 3.4 Å (Fig. 5c; Supplementary Fig. 6). β-sheet motifs, bulky side chains, and five N-glycans per protomer are visible in our map, supporting this resolution. The derived atomic model aligns with recombinant HA structures^34,35^. To our knowledge, this constitutes the highest-resolution cryo-ET reconstruction of on-virion HA to date and establishes HA as the smallest macromolecule thus far resolved to near-atomic detail using cryo-ET. This accomplishment relied on precise microscope calibrations before data acquisition, such as coma free alignment and dose rate optimization. These calibrations were continuously monitored through FlyTomo’s diagnostic feedback and intervened if deteriorating.

We also assessed FlyTomo on a complex residing inside the viral envelope. The VSV RNP forms a helical assembly of subunits, each comprising nine RNA nucleotides bound to an N protein that interacts with inner (IM) and outer (OM) matrix proteins. This M-M-RNP assembly is highly flexible, with variable helical parameters that complicate high-resolution analysis^36,37^. Earlier cryo-ET work attained 7.5 Å resolution for the VSV trunk^38^. From 122 tomograms, 147 virions were selected for analysis. Rigid-body alignment of trunks, followed by subboxing, yielded a 4.6 Å map with local resolution to 4.3 Å (Fig. 5d; Supplementary Fig. 7). The map agrees with the published atomic model (PDB:7UWS)^37^. RNA, N, IM, and OM densities are readily apparent at this resolution (Fig. 5d). Back-projection of subtomogram averages confirmed the continuous helical lattice and RNA density, consistent with prior observations^36,37^. Precise TS alignment and tomogram reconstruction are essential to prevent distortion of this helical lattice. FlyTomo’s integrated diagnostics provided rigorous quality control; together with Off-EM-Session inspection and curation, this workflow preserved data integrity and was critical for resolving the flexible RNP at high resolution.

Collectively, these benchmarks demonstrate FlyTomo’s capacity to handle large datasets of low-abundance POIs (HCoV-229E S), small macromolecules (IAV HA), and highly heterogeneous assemblies (VSV RNP). The results further indicate the potential of FlyTomo for studying viruses in cellular contexts, where heterogeneity and low copy numbers present additional challenges.

### Testing FlyTomo on cellular tomography samples

To further test FlyTomo on cellular samples, we evaluated its performance on cryo-FIB-milled cellular lamellae from *S. cerevisiae* (EMPIAR-11658). These lamellae, typically characterized by the absence of fiducial markers, increased thickness, and crowded intracellular environments, pose significant challenges for TS alignment due to their low SNR (Fig. 6a). Using FlyTomo’s automated AWI detection algorithm, we measured sample thickness across the dataset. TS alignment was performed via the embedded AreTomo module, showing that lamella thickness (*alignZ*) ranged from 100 to 350 nm (Fig. 6b). Based on the measured thickness values, the dataset was stratified into five groups for alignment refinement. Subsequently, 80S ribosomes were identified from 90 well-aligned thin tomograms and aligned to a 5.65 Å resolution structure using a final set of 53,228 particles (Fig. 6c, Supplementary Fig. 8, Supplementary Table 2). The result demonstrates FlyTomo’s effectiveness in handling challenging cellular tomography datasets.

**Fig. 6.**
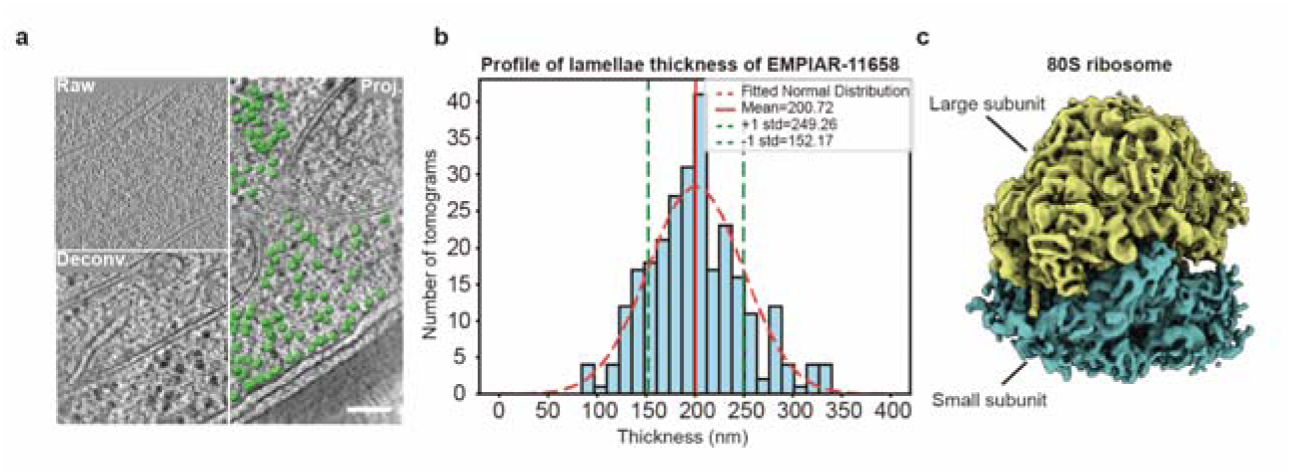
Performance of FlyTomo on cellular tomography sample. **a,** An exemplary slice of tomogram of S. cerevisiae lamella (EMPAIR-11658), showing raw (upper left), deconvolved (lower left), and ribosome-annotated tomograms (right). **b.** Histograms of pretilt and thickness of lamellae in EMPIAR-11658. **c** Final 80S ribosome structure at 5.65 Å from EMPIAR-11658.

In summary, our tests have demonstrated the advantages of FlyTomo in processing both molecular and cellular tomography samples. The high-throughput accelerates data processing; empirically validated parameters ensure data fidelity; integrated diagnostics support real-time decision-making on maintaining microscope alignment and excluding suboptimal data; and AWI-based geometry estimation enhances TS alignment accuracy. These advances highlight the power of integrated instrumentation and computation for high-resolution cryo-ET.

## Discussion

FlyTomo combines high-throughput data processing, real-time diagnosis and user-guided intervention for cryo-ET. Extensive validations across a diverse range of cryo-ET scenarios demonstrate that FlyTomo delivers reproducible results. For inexperienced users, FlyTomo’s empirically optimized default parameters and intuitive GUI reduce the learning curve of cryo-ET. For advanced users, FlyTomo has options for command line-based execution, a template file containing editable parameters, and a convenient offline intervention-reprocessing loop to flexibly refine their data. As an actively maintained platform, FlyTomo is positioned for iterative updates aligned with evolving user needs and methodological innovations.

FlyTomo has different design philosophy compared to the other streamlined cryo-ET data processing tools. Among the automatic data processing tools, current versions of Tomo Live^39^ and AreTomoLive^40^ only provide preprocessing pipeline until tomography reconstruction and denoising. Moreover, Tomo Live is a commercial plugin that is executable under Thermo Fisher Tomography 5. TomoBEAR^41^ provides a simplified and linear workflow to STA with limited feedback and has no GUI. Warp^42^ does not contain an automatic TS alignment function. nextPYP^43^, RELION5^25^ and ScipionTomo^44^ provide limited real-time diagnosis for microscope status and data quality during data acquisition. Other softwares, such as emClarity^45^, Dynamo, TOMOMAN^46^ and EMAN2^47^, are designed for standalone Off-EM-Session processing and require considerable cryo-ET experience to operate.

Nonetheless, the current version of FlyTomo has several limitations. (1) Its on-the-fly performance is contingent upon the speed and capacity of local data transfer and computing infrastructure, which may constrain its utility in cases of exceptionally high data volumes or limited computational resources; (2) Although the platform incorporates validated presets for ordinary cases, users working with unconventional specimen types may still require manual parameter tuning; (3) Its In-EM-Session module has not been extensively tested on cryo-lamellae data acquisition due to the lack of such samples in our laboratory.

FlyTomo will be smarter in the future development. By utilizing machine learning on large-scale datasets to identify intrinsic correlations between experimental conditions (e.g., sample characteristics, microscope stability and stage drift patterns) and results (e.g., data processing diagnostic metrics and final resolution), FlyTomo will develop signatures on specific samples and microscopes. Using the signature, FlyTomo could guide adaptive acquisition strategies tailored to individual setups. For instance, predictive models trained on historical datasets could recommend adjustments for microscope alignment, thereby improving the proportion of high-quality data. Furthermore, the signature will enable specialized parameters for data processing.

FlyTomo is well-suited for cryo-ET facilities aiming to enhance training, operational efficiency, and reproducibility; for research groups studying biological specimens that require immediate feature assessment; and for projects with limited microscope access and demand high structural yield from each session. As the field advances toward large-scale visual proteomics, software ecosystems that prioritize adaptability, automation, and usability will be indispensable to the future generation of structural biology.

## Materials and Methods

### Lamellae air-water interface detection

FlyTomo detects the AWI in tomograms by calculating Shannon entropy. To improve computational efficiency, the input tomograms were generated with a relatively high binning factor (e.g., bin32) using weighted back projection as the default reconstruction method. Initially, the raw tomograms *V* were processed using a 3D SIRT-like filter adapted from its 2D counterpart^48^, which can be expressed as follows:

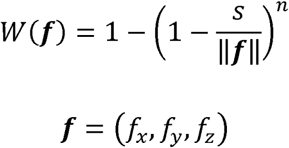

where *W* is the 3D SIRT-like filter, ||***f***|| is the norm of the 3D frequency vector ***f***, *s* and *n* are the step size and number of iterations of SIRT, respectively. Hence, we have the filtered tomogram *V_SIRT_* as:

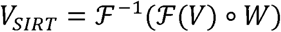

where *F*(.) denotes the Fourier transformation. Then, a new volume *E*, representing the randomness and complexity of *V_SIRT_*, is calculated, where each voxel value is computed by the Shannon entropy of the *k* x *k* neighborhood in the x-z slice of tomogram. To avoid the decrease of volume shape caused by the *k* x *k* sliding window, *V_SIRT_* is mirror-pad in advance. Therefore, we have:

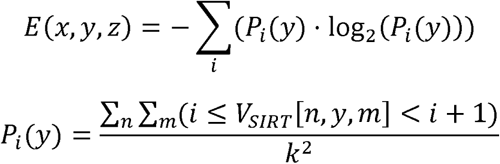

where *P_i_* (*y*) denotes the probability of the voxel (*x*, *y*, *z*) value within the *k* x *k* neighborhood falling in to range [*i*,*i* +1). For the purpose of acceleration, we expand the logarithm term in *E*(*x*, *y*, *z*) using a Taylor series up to the third-order term, and the *V_SIRT_* is in advance converted to the 8-bit grey scale volume. Therefore, we have:

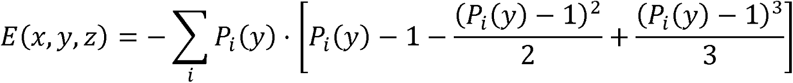

To label the lamella region, a segment threshold is calculated using Otsu’s method implemented in scikit-image, and the voxels where value higher than the threshold are marked as sample and the rest are background. The resulting segmentation is further processed with morphological closing in SciPy^49^ that smooths the surface of the labels. Hence, the final segmentation *S* is a 3D volume with the same shape of original tomogram, which labels the ice layer with ones and the air with zeros. Such 3D volume is thereafter converted to point clouds and subjected to RANSAC fitting in Scikit-learn^50^ to the general form of a plane equation:

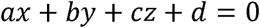

where *a*, *b*, *c*, *d* are constant parameters of the plane, which roughly describes the orientation and position of the lamella. We assume both of the AWIs are planes that are not necessarily parallel to each other. Therefore, the gradient of *S* is calculated, which ideally yields two flat layers of points on both sides corresponding to the upper and lower interface, followed by RANSAC fitting to two plane equations:

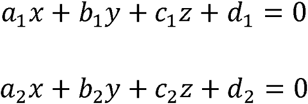

where subscript 1 and 2 denotes parameters of the upper and lower plane, respectively. Considering the uneven sample thickness throughout the whole lamella, the lamella thickness reported from FlyTomo is the thickness sampled from the center in x-y plane of the tomogram and modelled as a point-to-plane distance, namely, the distance from the sampling point on the upper surface to the plane of the lower surface. Therefore, we have:

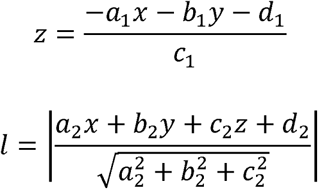

where *l* denotes the average sample thickness, (*x*, *y*, *z*) is the center coordinate of the sampling points on the upper surface. Likewise, the pretilt *θ* of the lamella is calculated as an average of the pretilt of upper and lower surface. Since the pretilt describes the angle offset along the y axis, we calculate it by computing the angle of the norm, projected to the x-z plane, of both the surface and x-y plane. Hence, we have:

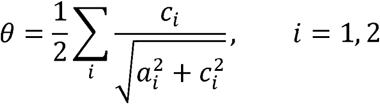

### Samples preparation and data processing of proteasome

The proteasome was prepared using a previously described method^51^. TS were acquired using a Titan Krios equipped with a Gatan K3 camera and GIF, with a slit width of 20□eV (Supplementary Table 2). Data was preprocessed in FlyTomo using marker-free TS alignment, and the resulting 8 × binned tomograms were imported to emClarity for template-matching (EMD-6278). The coordinates of the picked particles were imported into Dynamo for initial STA and refined by RELION4, including 3D refinement on 4 ×, 2 × and 1 × binning, followed by frame alignment and CTF refinement. A final 3.4 Å resolution map was resolved from 38,340 particles.

### Samples preparation, data acquisition and data processing of coronaviral S-trimer

A laboratory strain HCoV-229E virus (ATCC VR740) was obtained from the China Center for Type Culture Collection (CCTCC, Wuhan, China) and stored at −80 °C until use. Virions were propagated in Huh7 cells (Cell Resource Center of the Institute of Basic Medical Sciences, Beijing, China) for 5 days at 33 °C. The cell supernatant was collected and subjected to low-speed centrifugation (3,000 × g, 30 minutes, 4 °C) to remove cellular debris. For cryo-ET, discontinuous sucrose cushion was applied. Virions were sedimented through a 30% sucrose band to 50% band in phosphate-buffered saline (PBS, pH 7.4) by ultracentrifugation (Beckman SW32.1 rotor) at 100,000 g, 4 °C for 3 h. The virus band at the sucrose interface was collected and concentrated using an Amicon Ultra Centrifugal Filter (0.5 mL, 100 kDa cutoff; Millipore, Germany) at 4 □. The concentrated virions mixed with equal-volume 8% paraformaldehyde (final concentration 4%), and inactivated for 1.5 h at 4 °C. Virus propagation, purification, concentration and inactivation were carried out in a BSL-2 lab at Tsinghua University.

Inactivated virions were mixed with 10 nm BSA-gold beads (Aurion, The Netherlands) and 4 μL of the mixture was applied to glow-discharged Quantifoil R2/1 holey carbon copper grids (200 mesh) in a Vitrobot Mark IV (Thermo Fisher Scientific, Waltham, MA) at 100% humidity and 8 °C. Grids were blotted for 4.5 s, plunge-frozen in liquid ethane, and stored in liquid nitrogen until imaging.

TS for HCoV-229E were acquired using a Titan Krios equipped with a Gatan K3 camera and GIF, with a slit width of 20□eV (Supplementary Table 2). TS data were preprocessed using FlyTomo as described above except the TS alignment was conducted by fiducial tracking. A total of 818 3D-CTF corrected tomograms of the HCoV-229E sample were reconstructed with 260 tomograms from pre-test session and 558 from the formal test session. During the four-day session, we select the TS collected from the first two days for structure determination before the session ending, named “In-EM-Session” group containing 245 tomograms and the remaining 313 tomograms was grouped naming “Off-EM-Session”.

Particle picking was performed as described previously^32^. Briefly, S-trimers were identified on 8 × binned IsoNet corrected tomograms by ilastik. The coordinates of S-timers were calculated and transformed into Dynamo-readable format with initial orientations assigned based on vectors normal to the local envelope. Totally, 21,279 S-timers from pre-test session, 20,713 from In-EM-Session, and 26,948 from Off-EM-Session were extracted into 48 × 48 × 48 voxels from 8 × binned tomograms in Dynamo. For subtomograms from pre-test session, an averaged map generated by initial coordinates was used as the template for Dynamo alignment with C3 symmetry applied. The S-trimer map generated after alignment was low-passed to 35 Å and then used as a template for alignment of subtomograms from In-EM-Session and Off-EM-Session. Aligned coordinates were exported to RELION4 for 3D classification and refinement. After removing suboptimal particles, 15,979 particles from the In-EM-Session group and 50,451 particles from all datasets were further refined with iterations of Frame alignment, CTF refinement and 3D refinement. The final maps from the In-EM-Session and Off-EM-Session datasets achieved global resolutions of 5.63 Å and 3.90 Å, respectively. The maps were sharpened (validated by Guinier analysis) and estimated for their local resolutions in RELION4.

### Samples preparation, data acquisition and data processing of influenza A virus

The laboratory strain IAV (A/Puerto Rico/8/34/H1N1) was obtained from CCTCC and stored at −80 °C until use. The virus was inoculated in 10-day-old embryonated hens’ eggs and incubated for 72 h. Viral particles were pelleted from allantoic fluid through a 33% (w/v) sucrose cushion via ultracentrifugation (112,000 × g, 1.5 h, 4 °C; Beckman SW32 rotor). After aspirating the supernatant, the viral pellets were resuspended in PBS, pH 7.4). The IAV was further purified using a 10%-60% (w/v) sucrose gradient. The IAV band was extracted, dialyzed to remove sucrose, and concentrated using Amicon centrifugal filters (0.5 mL, 100 kDa cutoff; Millipore, Germany).

Purified IAV virions were mixed with 10 nm BSA-gold beads (Aurion, The Netherlands). 4 μL mixture was applied to glow-discharged Quantifoil R2/2 holey carbon copper grids (200 mesh) in a Vitrobot Mark IV (Thermo Fisher Scientific, Waltham, MA) at 90% humidity and 8 °C. Grids were blotted for 4-5 s, plunge-frozen in liquid ethane, and stored in liquid nitrogen until imaging.

TS for IAV were acquired using a Thermo Fisher Titan Krios G4 equipped with a Falcon4i camera and Selectris X, with a slit width of 20□eV. Data were collected using the BIS-TOMO data acquisition script developed by National Multimode Trans-scale Biomedical Imaging Center based on SerialEM (Supplementary Table 2). TS data were preprocessed using FlyTomo. All datasets underwent diagnosis within FlyTomo, during which dark frames were removed, and gold fiducial localization and distribution were optimized, thereby reducing the fiducial mean error to < 0.2 nm for the majority of the datasets. Following optimal alignment calibration, a total of 65 tomograms of IAV samples were reconstructed and 3D CTF corrected in FlyTomo, resulting in a final pixel size of 0.92 Å/pixel. For subsequent particle picking, the tomograms were 8 × binned, and missing wedge correction was applied using IsoNet. Membrane segmentation was then performed on the corrected tomograms using MemBrain-Seg. Normal vectors to the membrane surface were generated using custom laboratory scripts employing Poisson surface reconstruction and sampled by the intervals corresponding to half the HA head diameter (i.e., 4 nm, based on an HA head diameter of 8 nm). These normal vectors served as initial seeds for STA.

A total of 260,571 particles were extracted from the 8 × binned tomograms with a box size of 32³ voxels. For initial model generation, 17,058 particles were randomly selected from five representative tomograms. STA was conducted in Dynamo, utilizing the PDB:1RU7, low-pass filtered to 20 Å, as an initial template. The resulting average served as the initial model for a global Dynamo alignment encompassing all particles. After three iterative alignment rounds and removal of overlapping particles, a dataset of 118,489 particles was obtained. Subsequently, this refined average, along with PDB:8E6J (low-pass filtered to 20 Å), were employed as templates for multi-reference classification within Dynamo. This step facilitated the removal of poorly aligned particles, yielding a final set of 97,201 particles. These particles were then exported to RELION4 for multiple rounds of 3D refinement and frame alignment imposing C3 symmetry. A final map at 3.6 Å resolution was resolved. Local resolution was estimated using RELION4.

### Samples preparation, data acquisition and data processing of VSV

VSV Indiana serotype seeds, rescued^52^ by plasmids of pCAG-VSVL (Addgene plasmid # 64085), pCAG-VSVN (Addgene plasmid # 64087), pCAG-VSVP (Addgene plasmid # 64088), pVSV Venus VSV-G (Addgene plasmid # 36399)^53^ and pC-T7^54^ was used to infect BSR-T7/5 cells at a multiplicity of infection (MOI) of 0.001. After 24 h, the infected cell supernatants were collected and centrifuged to remove large impurities. The virus was then concentrated with a centrifugal filter (100 kDa MWCO, Millipore), and acidified by 10 mM HEPES, 140 mM NaCl buffer (pH 5.5) at 37 □ for 15 min. 3 μL purified virions were applied to a glow-discharged 200 mesh holey carbon film-coated copper grid (R2/1; Quantifoil, Jena, Germany) and blotted for 3.2 s before plunge-freezing into liquid ethane using a Cryoplunge 3 (Gatan, Pleasanton, CA).

A total of 122 TS were collected and processed using FlyTomo, with TS alignment using marker-free mode, followed by manual intervention (Supplementary Table 2). A total of 147 virions were manually selected from the reconstructed tomograms, with their orientations and positions determined by identifying the head and tail coordinates. To define the trunk region, midpoint of each coordinate pair was calculated as the central point for the trunk. The helical trunk region was further refined at 8 × binning with the box size of 40³ voxels using Dynamo. The helical parameters were defined as a rise of 1.34 Å per subunit, a helical turn of 9.35° (Supplementary Fig. 7).

Subboxes containing viral RNP subunits were re-extracted from the aligned trunk region at 8 × binning (pixel size 5.44 Å) based on these helical parameters, with a box size of 40³ voxels. To improve the accuracy of particle selection, we restricted subbox extraction to the central trunk region within a defined length (Supplementary Fig. 7). Additionally, to prevent overfitting due to duplication, we extracted one subbox per five subunits, maintaining the independence (Supplementary Fig. 7) of half-sets. Initial alignments were performed in Dynamo, followed by further refinements in RELION4. The final reconstruction achieved a global resolution of 4.6 Å, with a central resolution of 4.3 Å. With the particle orientations and positions determined, the refined asymmetric subbox were back-projected into the tomogram space to visualize the viral structure within its native context.

### Cryo-ET data processing of 80S ribosomes from *S. cerevisiae* lamellae dataset

The raw movie stacks and SerialEM MDOC files of EMPIAR-11658 were prepared for test. Data preprocessing was performed using the FlyTomo with TS alignment using marker-free alignment from AreTomo. Manual intervention was performed to remove poor tilt images and reprocessing of the selected TS alignment was conducted. Lamellae thickness was evaluated before the fine TS alignment. All data was divided into five groups based on 1 × binned thickness in pixels (1.96 □/pixel), with thresholds set at 500, 800, 1,100, and 1,400 pixels. In fine alignment, the *alignZ* for the groups were assigned as 300, 600, 900, 1,200, and 1,500, respectively. 90 sets of TS with samples thinner than 1,100 pixels (215.6 nm) were selected, reconstructed and 3D CTF-corrected by 4 × and 8 × binning in FlyTomo. The cytosolic 80S ribosomes were template matched on 8 × binned tomograms in FlyTomo using a yeast 80S ribosome map (EMD-0049) low-pass filtered to 40 □ as a template. The top 1,000 hits in a tomogram intersected with cross correlation over 0.25 were extracted, yielded a total of 87,422 particles for subsequent STA in Dynamo and RELION4.

Initially, 8,775 particles from 10 tomograms were cropped and averaged in 4 × binning in FlyTomo, followed by 3D classification and 3D refinement to 15.92 □ (n = 4,123) in 4 × binning using RELION4. In the following STA of all the particles from 90 tomograms, the coordinates and orientations from template matching were directly input to RELION4, utilizing the previous 4 × binned map as the initial refinement reference. False positives of template matching were excluded by the 3D classification. Particles were then refined consequently in 4 ×, 2 × and 1 × binned, and subjected to 3 iterations of CTF refinement and frame alignment in 1 × binning. All the masks in the above RELION4 processing were of ribosome shape. Finally, the 80S ribosome was resolved to 5.65 □ resolution from 53,228 particles.

## Acknowledgements

This work was supported in part from National Natural Science Foundation of China (#32241031, #82241066 and #32171195) and Tsinghua University Dushi Fund. We thank Dr. Jianlin Lei, Dr. Fan Yang and Dr. Xiaomin Li from the cryo-EM Facility, Technology Center for Protein Sciences, Tsinghua University, for their support on cryo-ET data collection. We thank the computational facility support on the cluster of Bio-Computing Platform (Tsinghua University Branch of China National Center for Protein Sciences Beijing). We thank Dr. Xiaojun Huang and Dr. Yan Zeng from National Multimode Trans-scale Biomedical Imaging Center for their support on cryo-ET data collection. We thank Dr. Xueming Li from Tsinghua University for providing the proteosome plasmids.

## Declarations

### Author Contributions

S.L. conceived and supervised the project. Z.Z. developed the software. C.P., W.Z., K.L., J.L., Y.S. and S.L. processed the data and gave feedback for software improvements. W.Z., K.L. and S.L. developed tools for the software. J.L., C.P., Y.C., J.Z. and S.L. provided the sample and collected the data. R.L. built the atomic models. Z.Z., W.Z., C.P., K.L. and S.L. wrote the manuscript. All authors critically revised the manuscript.

### Data and materials availability

Electron microscopy maps of HCoV-229E S, IAV HA, and VSV RNP have been deposited in the Electron Microscopy Data Bank under accession codes EMD-XXXX, EMD-XXXX, and EMD-XXXX. Atomic models of HCoV-229E S, and IAV HA have been deposited in the Protein Data Bank under accession codes PDB-XXXX and PDB-XXXX.

