## Supplementary Info for "FlyTomo: A streamlined software for on-the-fly cryo-ET data processing and diagnosis"

**Supplementary Information**

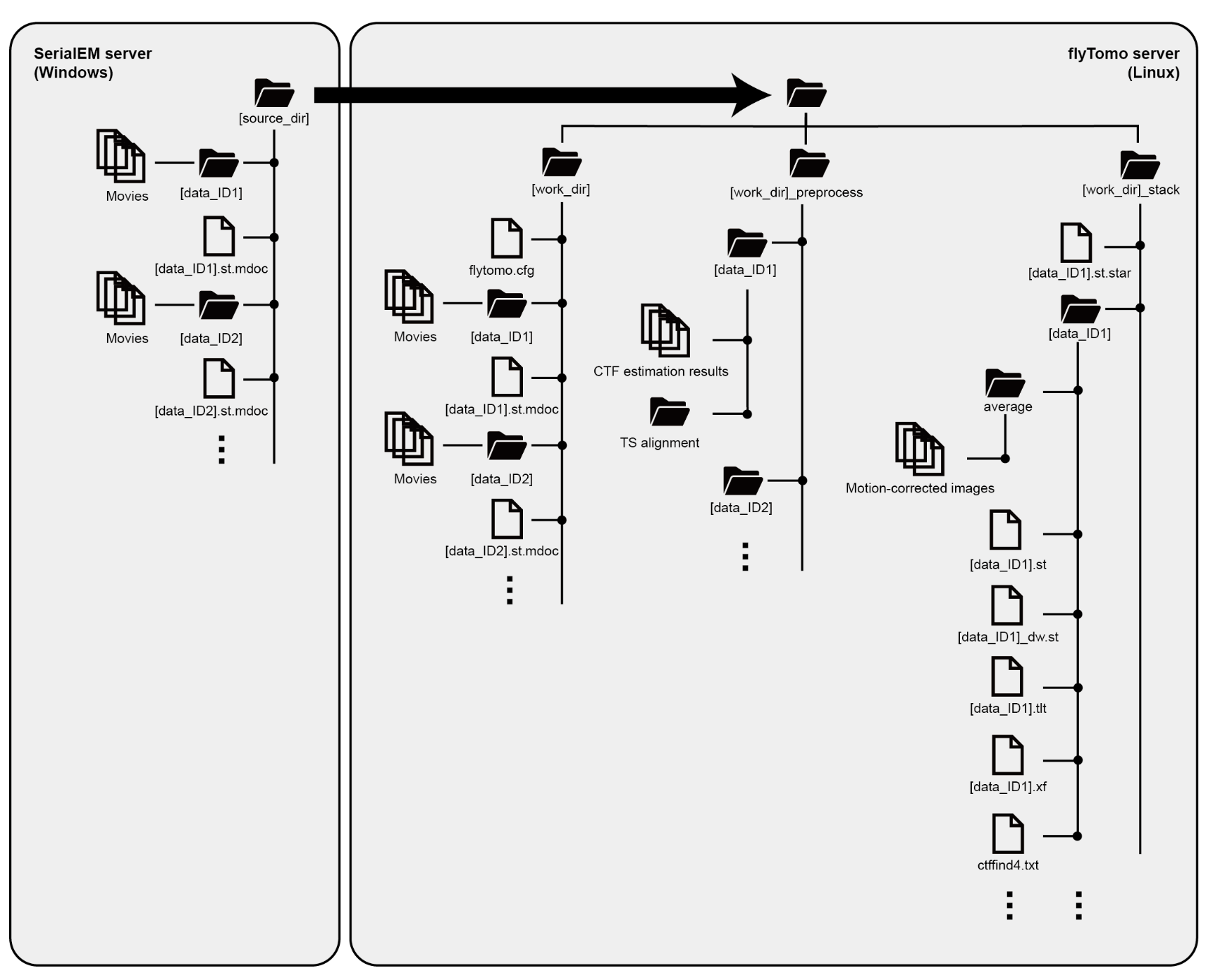

***Supplementary Fig. 1 | Folder organization of a FlyTomo project.*** *The raw movies and metadata are collected and stored in the [source_dir] on SerialEM server. During FlyTomo processing, data are transferred to the FlyTomo server and the results are categorized into [work_dir], [work_dir]_preprocess, and [work_dir]_stack folders.*

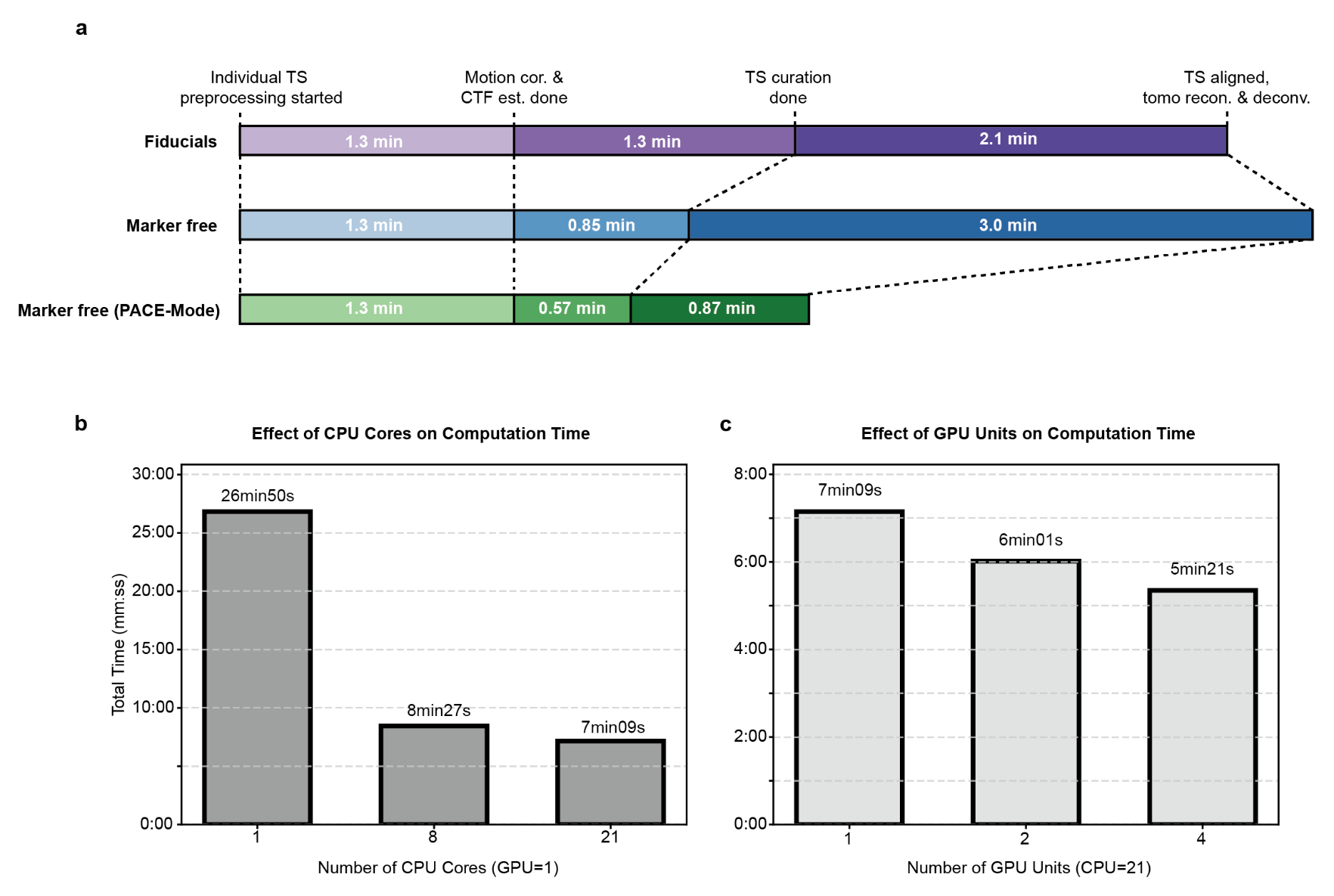

***Supplementary Fig. 2 | Benchmarking FlyTomo preprocessing time. a,*** *Time per tomogram for three preprocessing pipelines using different TS alignment methods: fiducial-based (purple), marker-free (blue), and marker-free with PACE-mode enabled (green).* ***b,*** *Preprocessing time (marker free) by different number of CPU cores, with GPU count fixed at one.* ***c,*** *Preprocessing time (marker free) by different number of GPU units, with CPU count fixed at 21.*

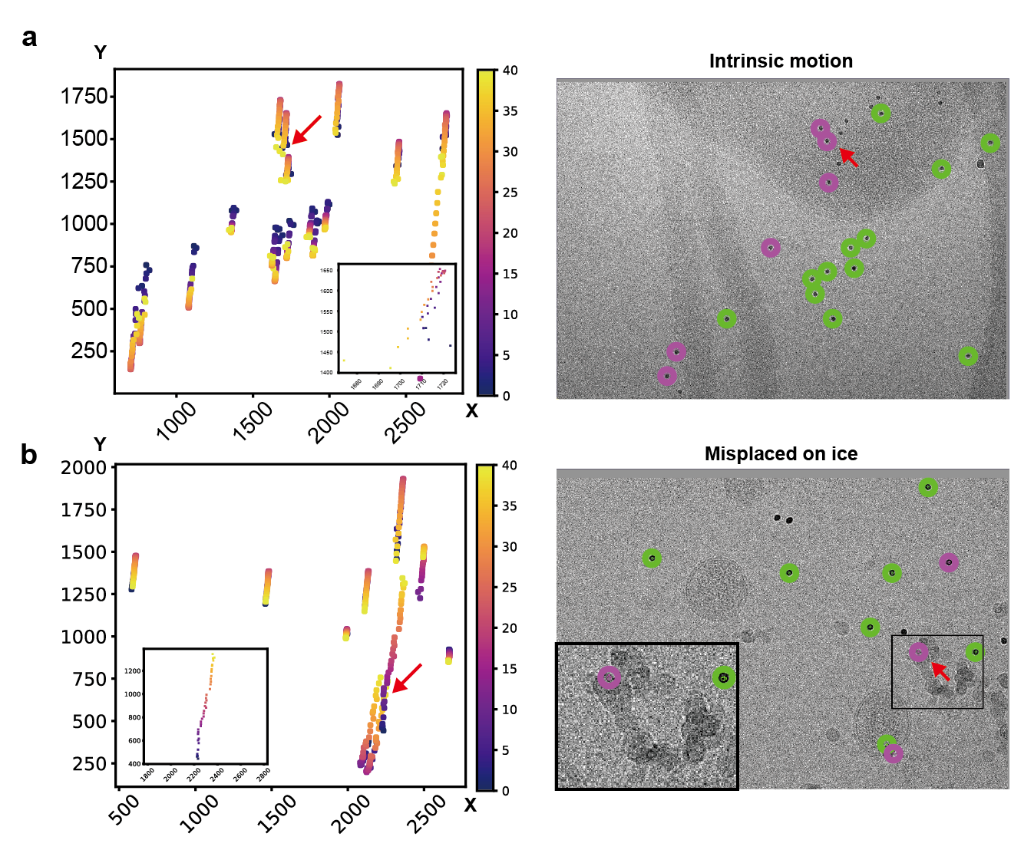

***Supplementary Fig. 3 | Additional feedback on data quality and processing performance.*** *Trajectory plots visualize fiducial movement across TS.* ***a****, Example of intrinsic sample motion: a single fiducial exhibits stray displacement (red arrow, inset) while others remain stable.* ***b,*** *Example of a misplaced fiducial: erratic trajectory and positional jump (red arrow, inset). Micrograph confirms its location on a contaminant ice crystallite outside the AOI, making it unsuitable for accurate alignment. Tilt numbers are color-coded.*

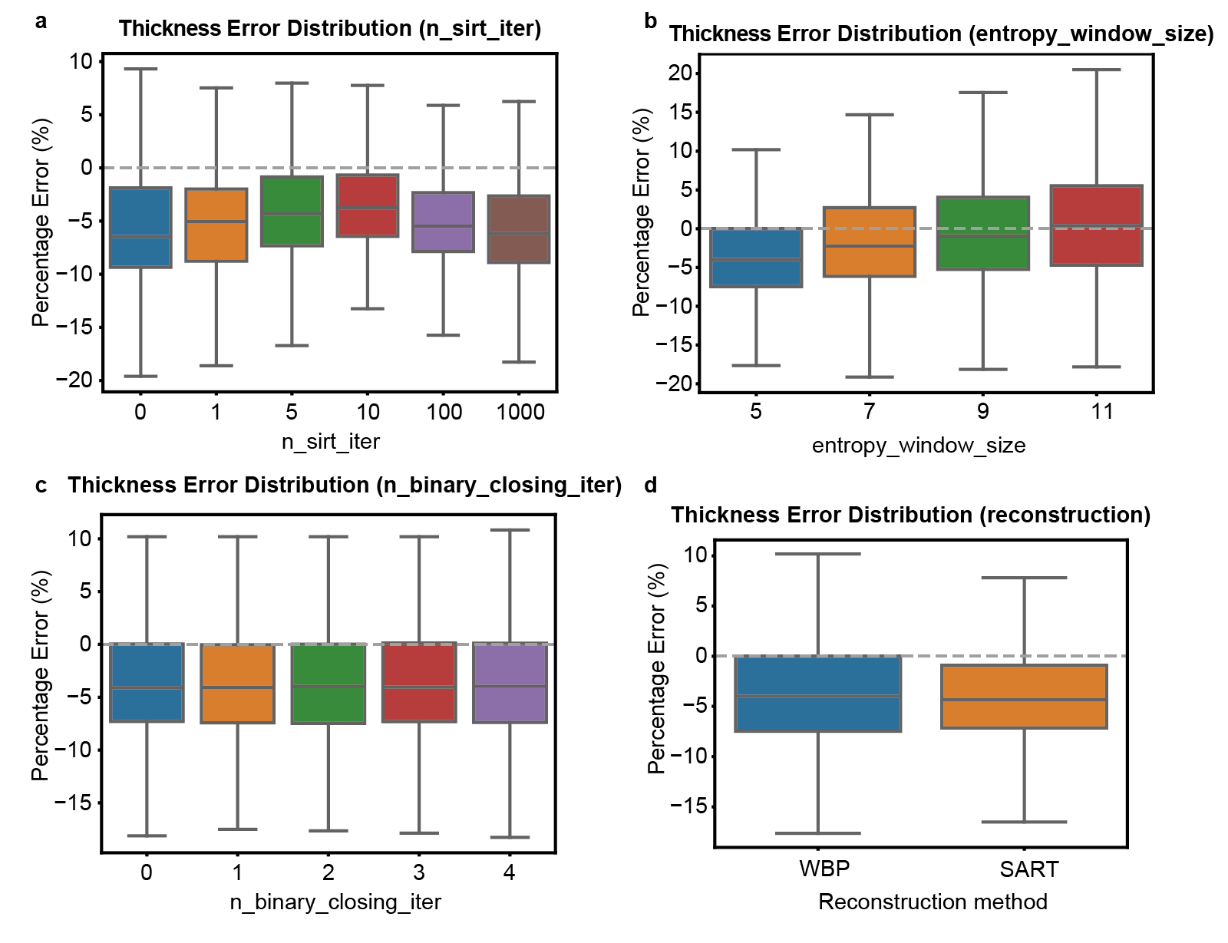

***Supplementary Fig. 4 | Tuning the parameters of AWI detection algorithm. a-d****, Distribution of percentage error in thickness estimation with parameters tuned one at a time, including n_sirt_iter (****a****), entropy_window_size (****b****), n_binary_closing_iter (****c****), reconstruction method (****d****).*

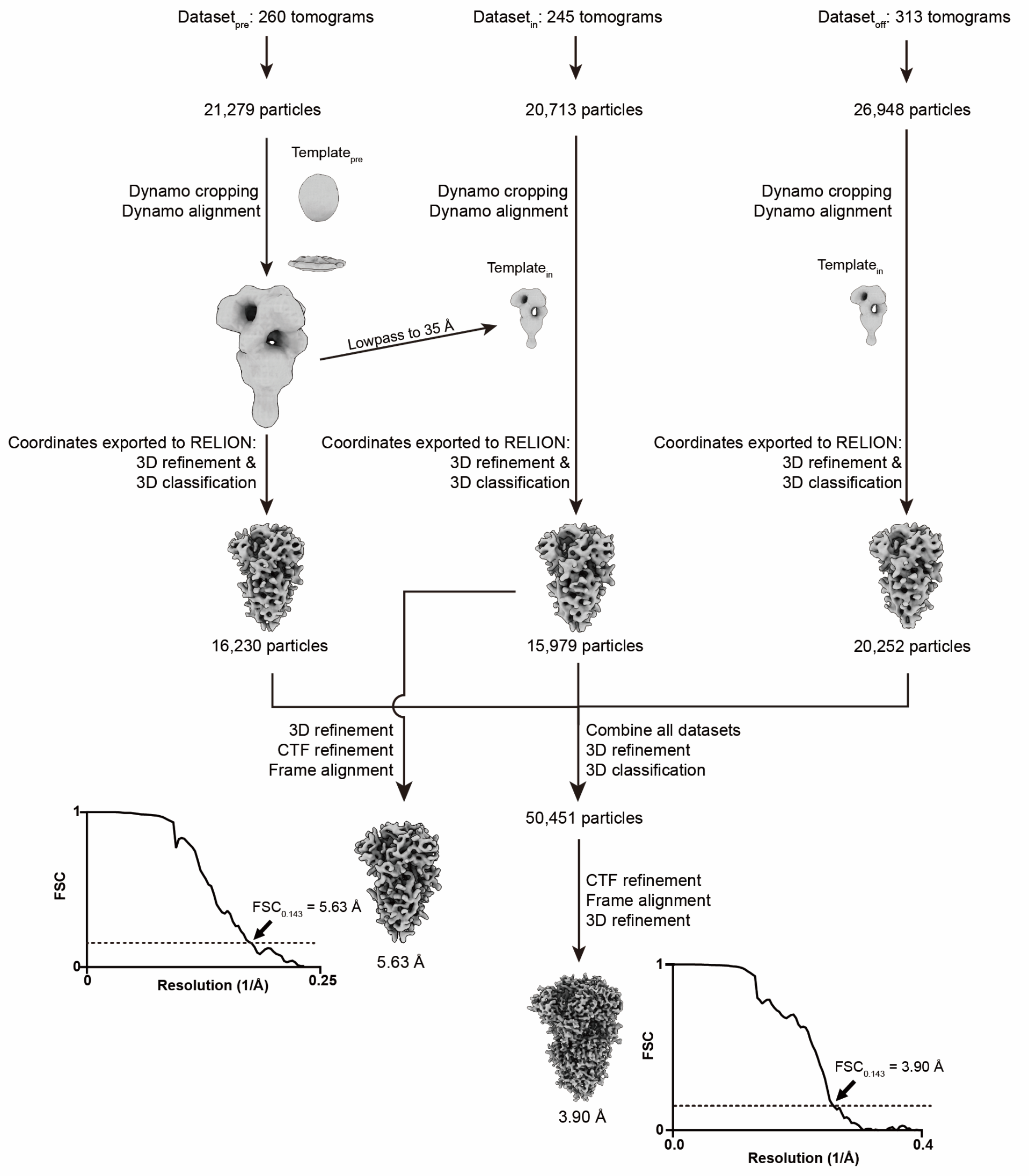

***Supplementary Fig. 5 | STA for HCoV-229E S trimer.*** *A total of 818 tomograms were collected across three parts (Dataset_pre_, Dataset_in_, and Dataset_off_). and cropped in Dynamo. Template generated de novo from Dataset_pre_ were used for particle alignment across all three datasets. After initial processing and 3D classification in RELION, an intermediate map with a resolution of 5.63 Å was generated by combining the clean particles from the first two datasets.* *For the final high-resolution structure, the selected particles from all three datasets were merged (n = 50,451), subjected to a final round of 3D refinement, CTF refinement, and per-particle frame alignment in RELION, yielding a final map with a global resolution of 3.90 Å.*

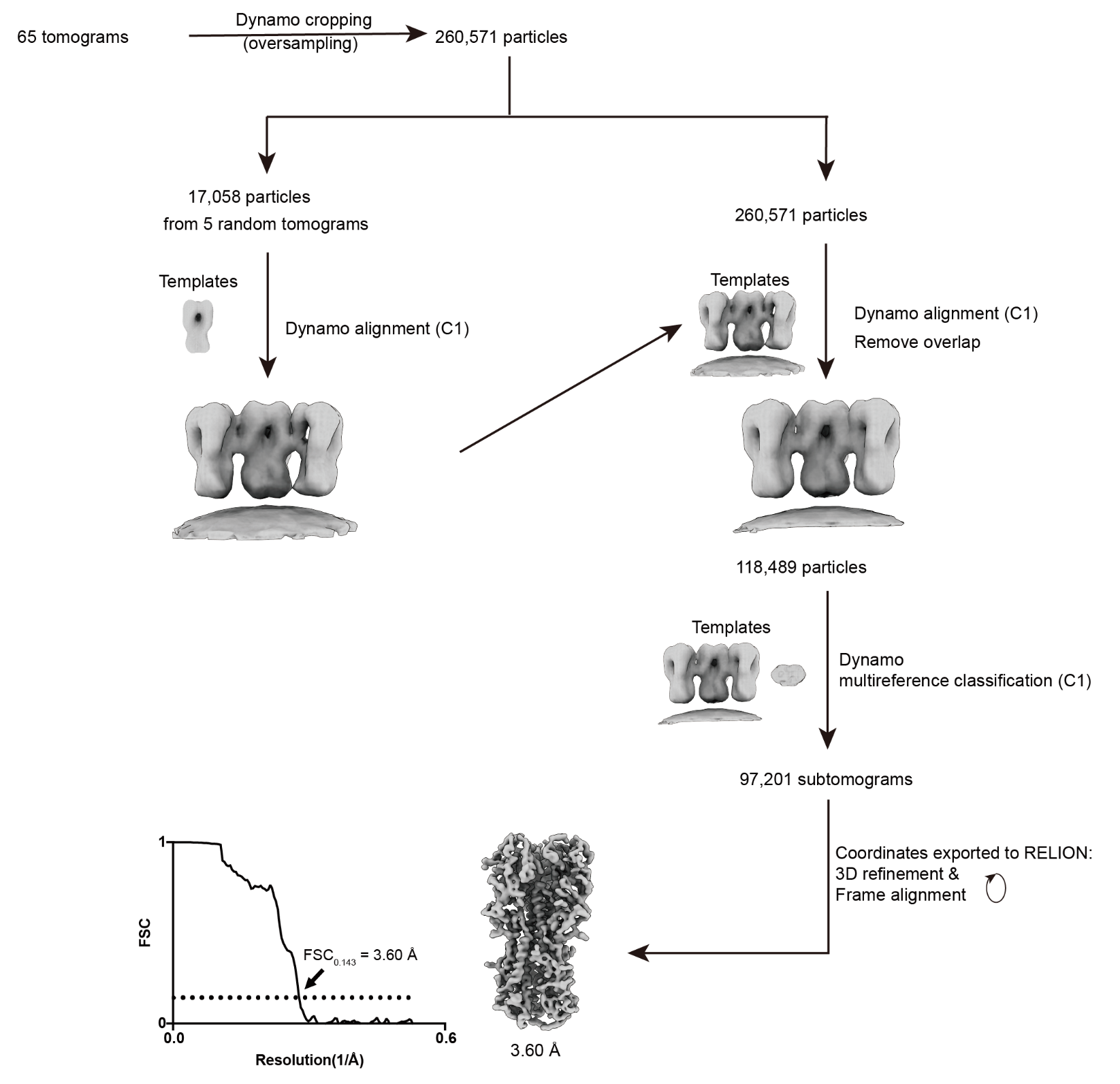

***Supplementary Fig. 6 | STA for IAV HA trimer.*** *An initial set of 260,571 particles was extracted from 65 tomograms using oversampling in Dynamo. A low-resolution template was generated by aligning a small subset of 17,058 particles from five representative tomograms (left path), and then used to align the full dataset. After removing overlapping particles, a set of 118,489 particles was obtained. This improved average was subsequently used as a reference for multireference classification in Dynamo to select for the most structurally homogeneous particles, yielding a final set of 97,201 subtomograms exported to RELION for high-resolution 3D refinement and Frame alignment. The final reconstruction reached a global resolution of 3.60 Å.*

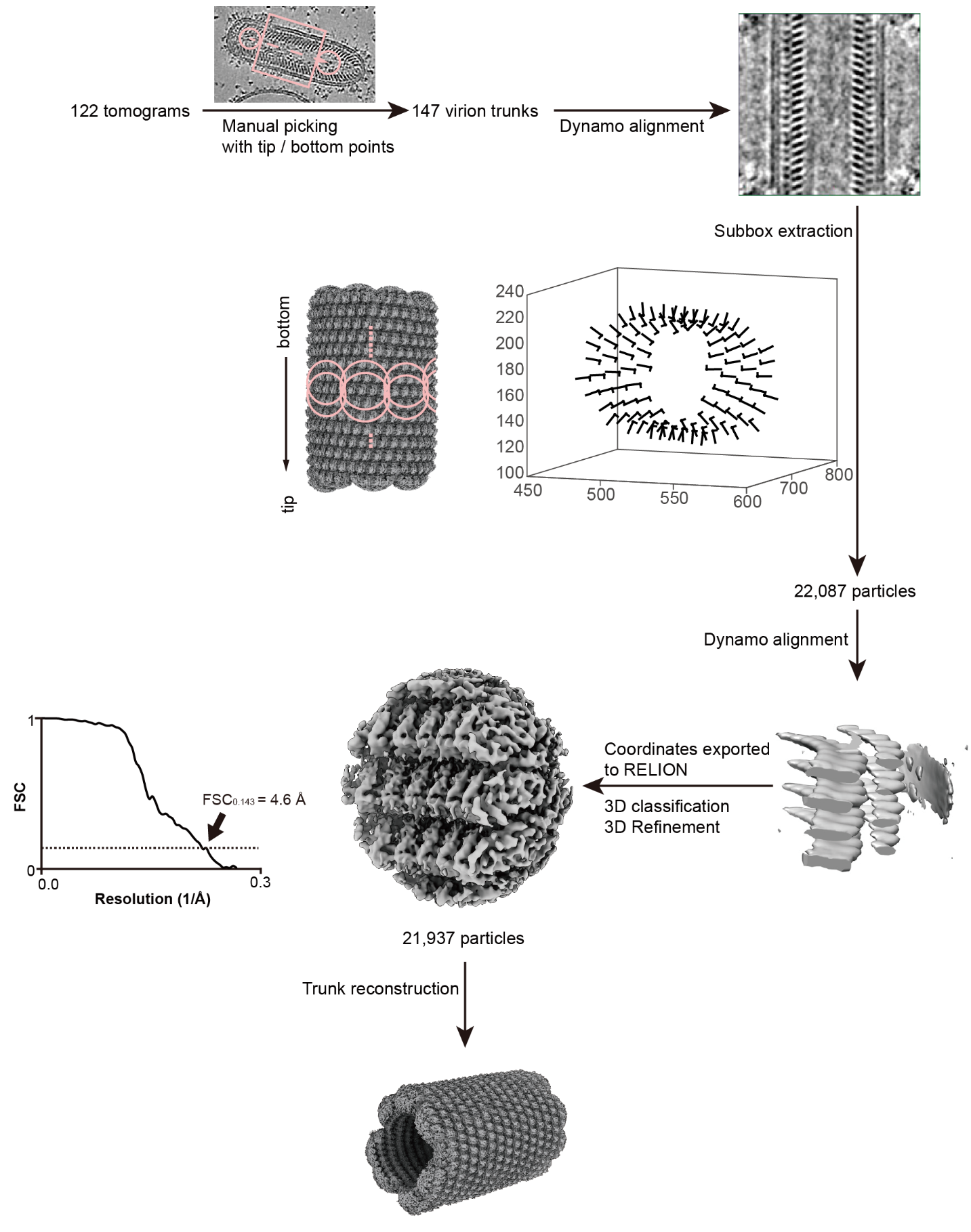

***Supplementary Fig. 7 | STA for VSV trunk using subboxing strategy.*** *From 122 tomograms, 147 VSV virion trunks were aligned as rigid bodies in Dynamo. Overlapping subboxes (n = 22,087) were extracted from trunks to address helical variation and refined separately. Final reconstruction from 21,937 particles reached 4.6 Å resolution (FSC 0.143 criterion). A composite trunk model was generated by projecting the refined subbox maps into tomogram space.*

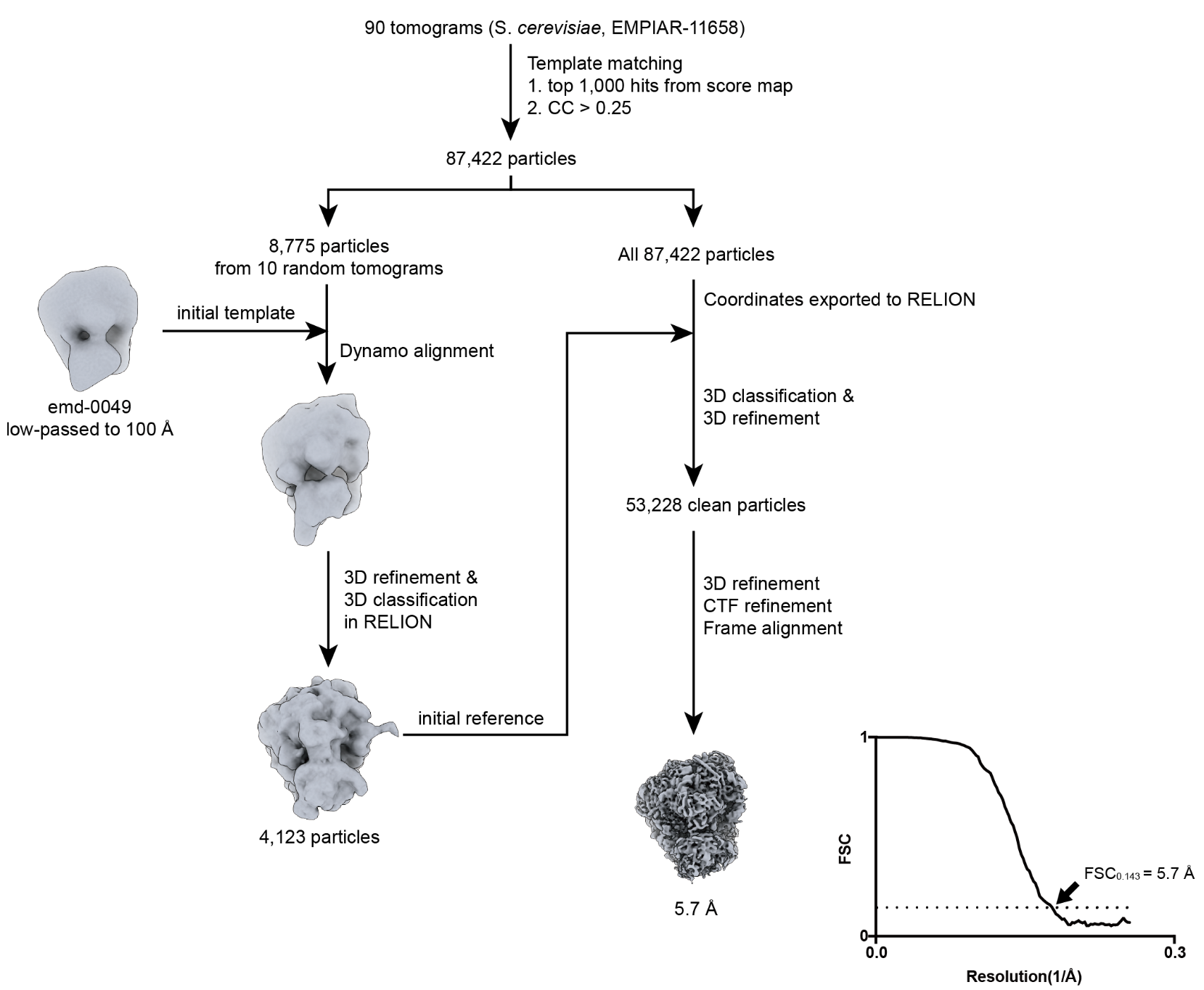

***Supplementary Fig. 8 | STA for 80S ribosomes in S. cerevisiae lamellae (EMPIAR-11658).*** *From 90 tomograms, 87,422 ribosomes were picked using template matching with an external map (EMD-0049). An initial reference was generated in Dynamo and used for RELION-based classification, yielding 53,228 particles. Final 3D refinement, CTF refinement, and per-particle frame alignment produced a 5.7 Å map (FSC 0.143 criterion).*

**Supplementary Table 1 | I/O reduction rate by the data compression and in-memory computation modules in FlyTomo.**

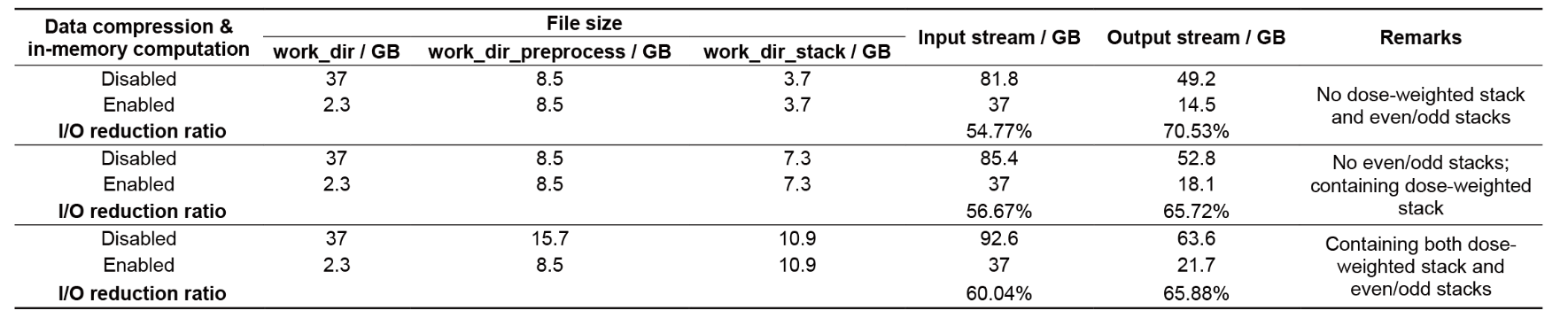

**Supplementary Table 2 | Cryo-ET data collection and subtomogram averaging statistics.**

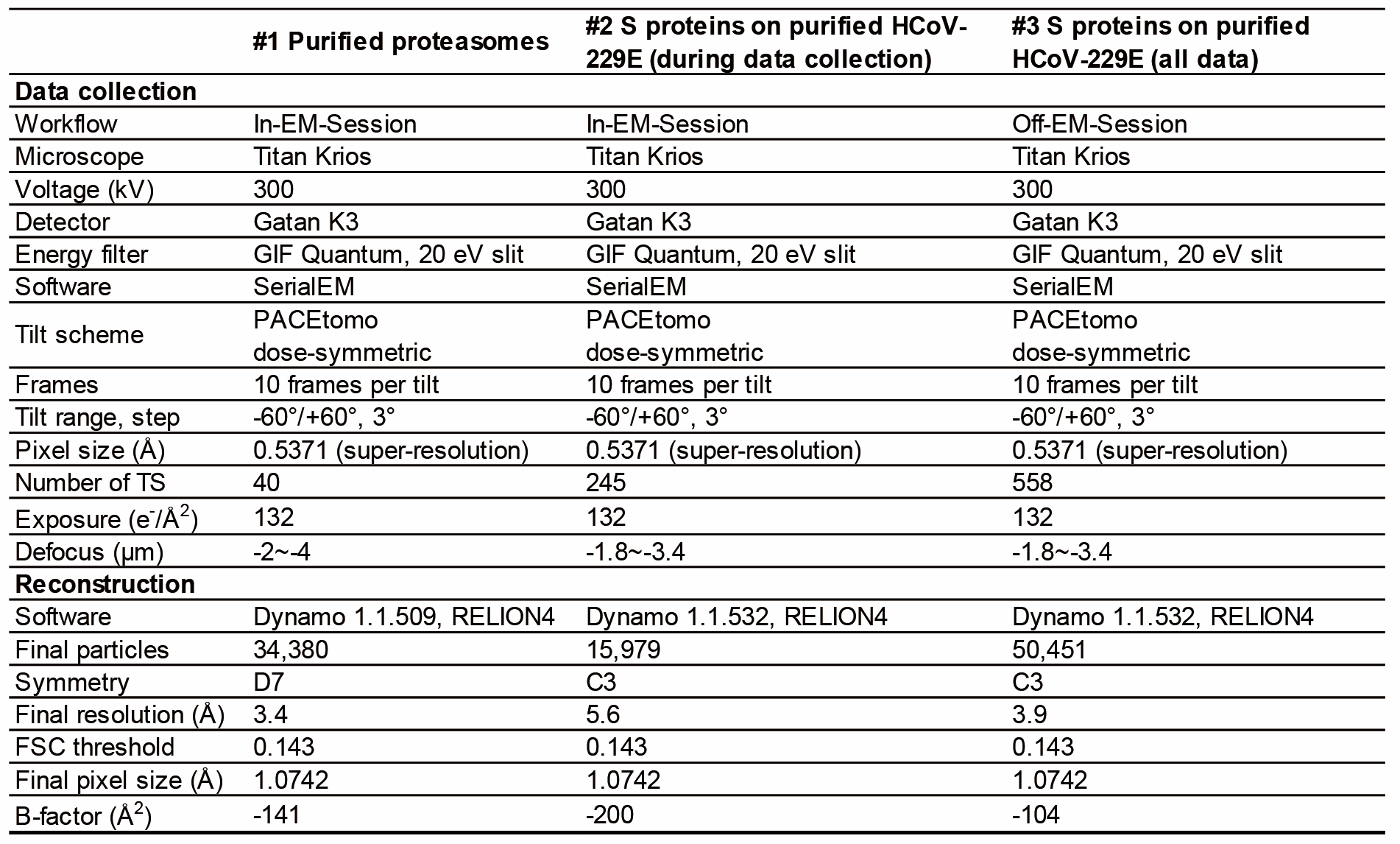

**Supplementary Table 2 continued | Cryo-ET data collection and subtomogram averaging statistics.**

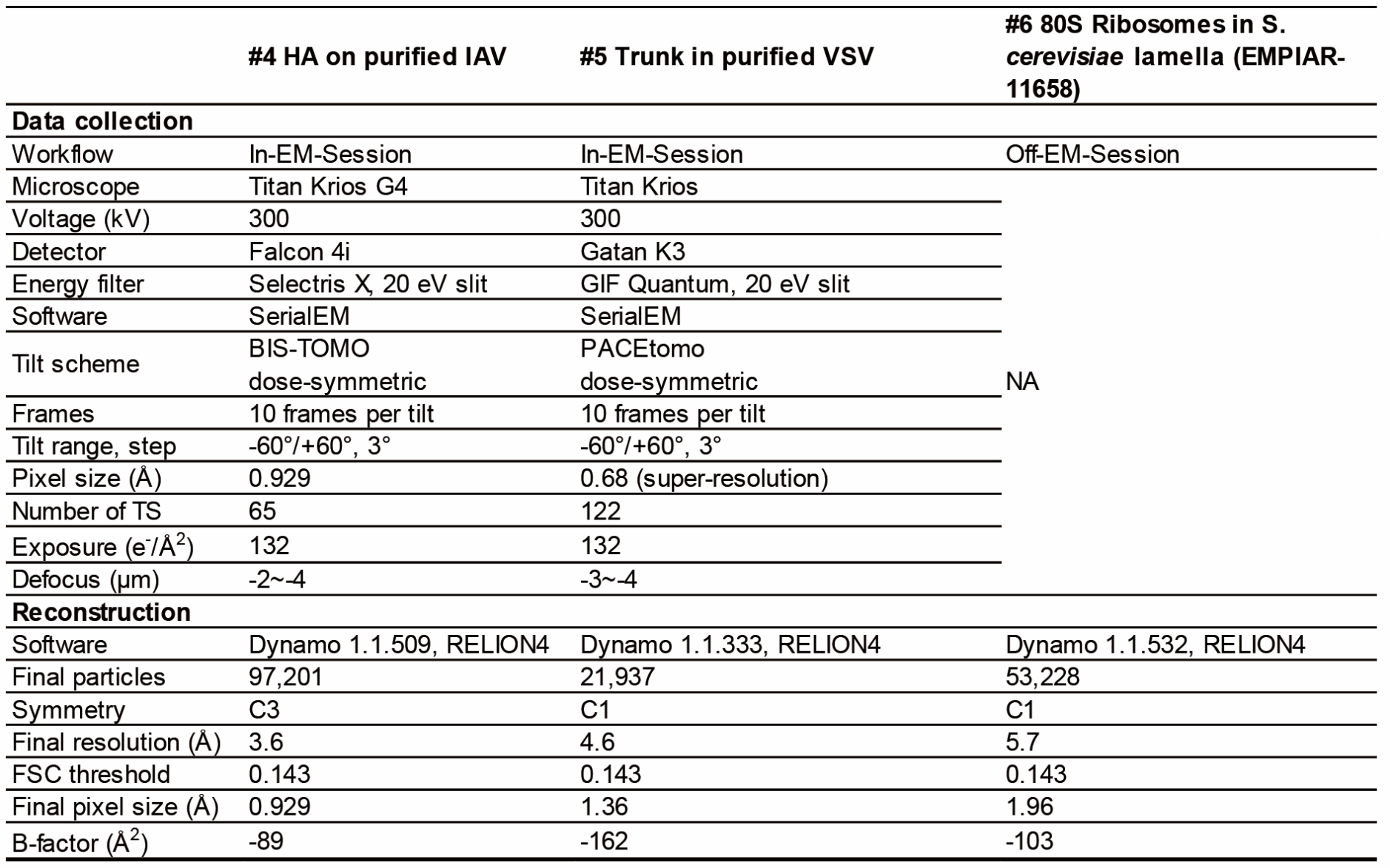
